# Mapping Natural Product Biosynthetic Hotspots: Prioritizing Conservation for Medicinal Resources

**DOI:** 10.1101/2023.08.12.553062

**Authors:** Muhamad Fahmi, Kojiro Takanashi, Yusuke Kakei, Yasuhiro Kubota, Keiichiro Kanemoto

**Affiliations:** Research Institute for Humanity and Nature, 457-4 Motoyama, Kamigamo, Kita-ku, Kyoto, 603-8047, Kyoto, Japan; Department of Science, Graduate School of Science and Technology, Shinshu University, 3-1-1 Asahi, Matsumoto, 390-8621, Nagano, Japan; Department of Biology, Faculty of Science, Shinshu University, 3-1-1 Asahi, Matsumoto, 390-8621, Nagano, Japan; The Institute of Vegetable and Floriculture Science, National Agriculture and Food Research Organization, 3-1-1 Kannondai, Tsukuba, 244-0813, Ibaraki, Japan; Faculty of Science, University of the Ryukyus, 1, Sembaru, Nishihara, 903-0213, Okinawa, Japan; Think Nature Inc., Nishihara, 903-0213, Okinawa, Japan; Graduate School of Environmental Studies, Tohoku University, Aoba, 468-1, Aramaki, Aoba-ku, Sendai, 980-8572, Miyagi, Japan; Research Center for Social Systems, Shinshu University, 3-1-1 Asahi, Matsumoto, 390-8621, Nagano, Japan

## Abstract

**Background:** Natural products (NPs) are vital for promoting human health. Given the increasing threats of biodiversity loss and ecosystem degradation, understanding the geographical hotspots of NPs is essential to strategically prioritize areas for conservation, ensuring the sustained availability of these invaluable medicinal resources.

**Methods:** We constructed a global diversity map for 1,434 woody angiosperm species, each represented by existing genomic or transcriptomic data. We curated a list of 166 enzymes essential for the biosynthesis and structural diversity of NPs, and identified geographical hotspots of NPs by averaging enzyme presence across the grids. We also examined the distribution pattern of each enzyme. To gain deeper insights into NP distribution patterns, we performed a comparative analysis of enzyme groups responsible for the biosynthesis of two pharmacologically significant compounds with distinct biosynthetic pathways, shikonin derivatives and benzylisoquinoline alkaloids.

**Findings:** Our study reveals a correlation between NP hotspots and biodiversity hotspots, with a subset of markers displaying unique, region-specific patterns. Comparative analysis of enzymes for shikonin derivatives and benzylisoquinoline alkaloids biosynthetic pathways shows similar pattern, with the former demonstrating a unique and region-specific distribution.

**Interpretation:** Our findings emphasize the importance of preserving biodiversity hotspots for sustaining NP-based medicinal resources. Additionally, the specific distribution of certain enzyme markers, such as those related to shikonin derivatives, suggests that some NPs may necessitate targeted conservation strategies. This study provides a foundational roadmap for identifying the geographical hotspots of NPs and developing targeted conservation strategies.

**Funding:** Japan Society for the Promotion of Science through the Grant-in-Aid for Scientific Research (A) 20H00651.

## Introduction

Natural products (NPs), especially those derived from plants, have served as the cornerstone of human medicinal practices for millennia.^1^ The earliest evidence of such use dates back to Mesopotamia, approximately 2600 BCE.^2^ NPs are valuable to the pharmaceutical and biotechnology industries, which have produced numerous modern medicines from them. Indeed, by the middle of the 1970s, NPs were the main source of new therapies in the pharmaceutical drug pipeline. Between the 1980s and the 2010s, unmodified NPs (5%), NP analogues (28%), and NP pharmacophores (35%) accounted for two-thirds of all modern medications.^3^ Currently, the pharmaceutical landscape leans more towards semisynthetic and fully synthetic drugs, reducing their reliance on NPs.^2^ Despite this shift, the genetic resources integral to NP biosynthesis continue to hold substantial value for the development of medical therapies and treatments. These genetic resources, consisting of enzymes and biosynthetic pathways involved in the formation of NPs, are key tools that allow us to explore, recreate, and improve natural chemical structures. Even for synthetic drugs, knowledge derived from the study of NP biosynthesis can guide the design of new molecules and the development of more efficient production methods.^4^

A multitude of genetic resources play an indispensable role in shaping the future of healthcare and advancing the development of medicinal applications. For instance, the cytochrome P450 monooxygenase family (CYP enzymes) stands out for its remarkable versatility and breadth. These heme-containing proteins catalyze a wide range of oxidation reactions during the postassembly phase of NPs.^5^ The importance of CYP enzymes is exemplified by CYP725A4, which is involved in the biosynthesis of Taxol, a potent anticancer agent derived from the Pacific Yew tree (*Taxus brevifolia*).^6^ Similarly, CYP71AV1 contributes to the biosynthesis of artemisinin, an effective anti-malarial drug from the sweet wormwood plant (*Artemisia annua*).^7^ Beyond their crucial roles in natural biosynthesis, these enzymes have been harnessed for metabolic engineering, utilizing heterologous microorganisms for the renewable production of drugs.^8^ Importantly, these examples reiterate the importance of protecting biodiversity, as it harbors genetic resources vital for the development of medicine.

The amount and quality of genetic resources for medicine depend heavily on functioning natural ecosystems.^9^ Three decades of research have demonstrated that biodiversity loss can impede such functioning, and the current dominance of humans over global ecosystems has resulted in a significant rise in the rate of extinction.^9^ Consistently, assessments by the International Union for Conservation of Nature (IUCN) Red List revealed that over 41,000 species are currently at risk of extinction.^10^ Moreover, according to one study, between 2000 and 2010, 5.96% of worldwide forest cover was lost.^11^ Deforestation was particularly prevalent in tropical regions, with South American rainforests experiencing nearly half of the loss. Another study found that 27% of the loss was due to commodity-driven deforestation, with steady rates shifting geographically from Brazil to other tropical forests in Latin America and Southeast Asia.^12^

Given the significant threat to global biodiversity and the constraints on conservation resources, there is an imperative need for urgent and targeted conservation efforts. However, before such actions are taken, an understanding of the present state of biodiversity, the likely response of biodiversity to global environmental change, and the best conservation practices per region are needed. Many studies have already investigated species richness from local to global dimensions and recent approaches have included metrics to evaluate how mitigating threats and restoring habitats in specific locations can contribute to global biodiversity objectives.^13–17^ Furthermore, to enhance conservation strategies, genetic diversity assessments, mostly using mitochondrial markers, have been combined with phylogenetic and functional diversity to better understand the adaptability and evolutionary potential of species and ecosystems.^18–21^ However, the connection between these biodiversity patterns and the current state of NPs, especially those with pharmaceutical potential, remains underexplored. This study seeks to bridge this gap by mapping the geographical distribution of key biodiversity indicators linked to NPs with the potential for pharmaceutical application. By aligning conservation priorities with the untapped medicinal resources inherent in biodiversity, this work aims to guide not only the preservation of biodiversity but also the path toward the development of future medicines.

## Methods

### Study design

We conducted a genomic-driven, geospatial analysis in this study (Fig. 1). We collected genome and transcriptome data, as well as species distribution of woody angiosperms, from public databases and references (appendix 1 and 2). For species with only raw transcriptome data, de novo assembly was conducted. Given the intricate network of NPs biosynthesis, we curated a diverse array of enzymes crucial to the NPs biosynthesis and structural diversity as markers. This encompasses enzyme families such as CYP enzymes, which catalyze a wide range of oxidation reactions during the post-assembly phase of NPs, and UDP-glycosyltransferases (UGTs) which modify NPs by attaching sugar moieties. Other enzyme families, including O-methyltransferases (OMT), prenyltransferase (PT), DOXC-class 2-oxoglutarate-dependent dioxygenase (DOXC), flavin-containing monooxygenase (FMO), terpenoid synthases (TPS), BAHD acyltransferases (BAHD), and polyphenol oxidases (PPO), were also chosen due to their significant contribution to the NPs biosynthesis and differentiation. We curated 166 enzymes crucial for NP biosynthesis and annotated them in the genome and transcriptome assemblies, passing a completeness assessment (appendix 3 pp 4–20). A global species diversity map was created for 1,434 woody angiosperm species that had annotated enzymes, with a grid resolution of 110 km equal-area grid cells, ∼1 degree at the Equator (appendix 2). We estimated the richness of NPs by interpolating the average of key enzymes presence within each grid. Furthermore, we conducted a detailed distribution analysis for each individual enzyme. This allowed us to identify any enzyme with a unique distribution pattern that deviated from the global trends and might indicate unique regional biosynthetic capabilities or evolutionary histories. Complementing the broad-scale enzyme distribution analysis, we conducted a targeted investigation of two specific enzyme groups corresponding to the biosynthesis of shikonin derivatives and benzylisoquinoline alkaloids (appendix 3 p 21). These groups were selected due to their high pharmacological significance and distinctive biosynthetic pathways.^22,23^ By focusing on well-known, specific groups of enzymes responsible for the biosynthesis of two distinct NPs, we aimed to distinguish subtler details that may be overlooked when analysing aggregate enzyme markers and individual enzyme averages. We mapped the distribution patterns of these enzyme groups to determine whether their pathways exhibited biogeographical distributions that differed from those of the broader set of enzymes.

**Fig. 1.**
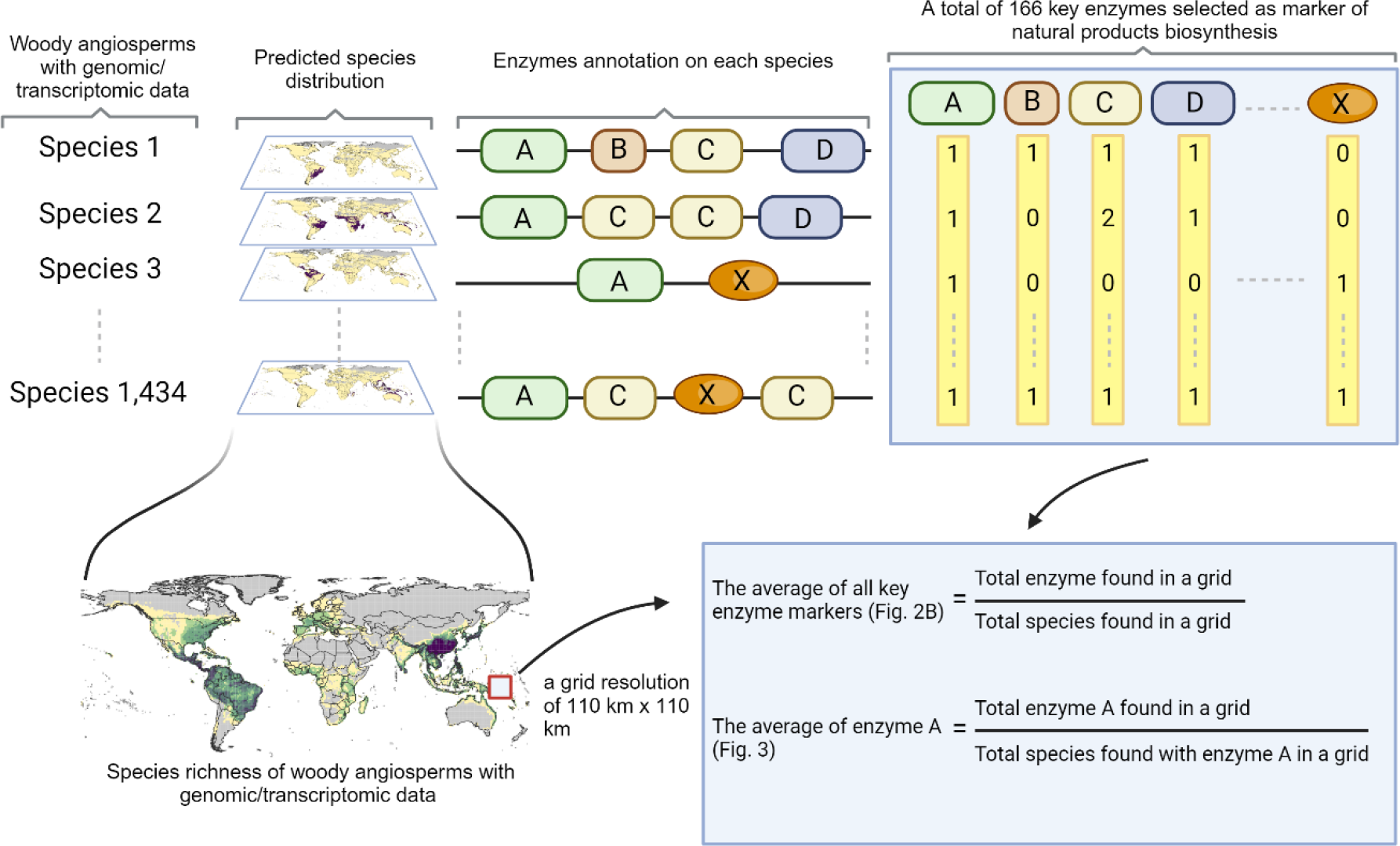
Illustration of genomic-driven, geospatial analysis. The process starts with the identification of woody angiosperm with genome or transcriptome data available in public databases and their predicted distribution on global maps. Each species is annotated with curated 166 key enzymes selected as markers for natural products biosynthesis. The lower part of the figure presents the species richness of woody angiosperms based on the genomic-transcriptomic data across a global map, with a 110 × 110 km grid cell. The calculations detail the average of enzyme markers and the proportion of specific enzymes found within the grid regions.

### Sequence Data Collection

We utilized the woody angiosperm species list outlined by ^24^ as a basis for determining a list of species. Their dataset is based on information in the national floras, including tree distribution checklists for each country, botanical literature, and various databases. We then curated the available genome or transcriptome data of each species from publicly accessible databases. The data incorporated assembled genomes, assembled transcriptomes, and raw transcriptomes. For each species, a single data type was selected based on availability, with a preference hierarchy of assembled genomes (most preferred), followed by assembled transcriptomes and raw transcriptomes (least preferred). The assembled genome data were collected from EnsemblePlants (release 52) and RefSeq, whereas the assembled transcriptomes were obtained from OneKP and Transcriptome Shotgun Assembly.^25^ For species lacking assembled genetic data, we collected their raw transcriptome data from the Sequence Reads Archive. To reduce the complexity in the assembly process of raw transcriptome data, third-generation sequencing data were omitted, and Illumina-derived data were preferred over data acquired from other second-generation sequencing instruments due to the abundance of the data. Additionally, only a single raw RNA-seq dataset from each experiment accession was collected for each species, and the data with the largest size were preferred. We collected 52 assembled genomes, 349 assembled transcriptomes, and 1211 raw transcriptome data for 1,612 species (appendix 1).

### De Novo Assembly for Raw Transcriptome Data

We performed de novo assembly by initially trimming the detected adapters, bases under 20 Phred score, and PolyG artifacts with a minimum length of 10 and by discarding reads with a read length equal to or below 36 after trimming using Fastp v.0.21.0.^26^ Then, multiQC v.1.10.1 was used to look at the results of the quality control of the RNA-seq data with similar read lengths.^27^ We removed the ribosomal RNA fragments using the SortMeRNA v.4.3.3 tool with SMR v4.3 default as the reference.^28^ We then performed de novo assembly at the scaffold level using SOAP denovo-Trans v.1.03 with the parameters set as follows: avg ins = 200, reverse seq = 0, asm flags = 3, and multiple k-mer lengths, including 21, 31, 35, and 41 (Assembly 1-4).^29^ For RNA-seq data with paired-end reads, before the de novo assembly process at the scaffold level, we first estimated the average insert size with the following steps. Contigs were assembled from RNAseq reads using SOAP denovo-Trans with parameters set as follows: avg ins = 200, reverse seq = 0, and asm flags = 3. Then, the first 400,000 RNA-seq reads were sampled and mapped to contigs with BWA-MEM, and the average insert size from the alignment was calculated. We later assembled RNA-seq contigs into scaffold levels using SOAP denovo-Trans v.1.03 with the estimated average insert size and the specified multiple k-mer lengths as above. We discarded scaffolds or contigs shorter than 200 bp and developed merged assemblies from the merging of assemblies 1-4 by cd-hit-est v4.6.1 with similarity thresholds of 93% (Assembly 5). We used assembly 5 for the subsequent analysis as this resulted in the best benchmark in the denovo assembly simulation (appendix 3 p 42; appendix 4). The pipeline for this de novo assembly was developed using Nextflow.^30^ Furthermore, we removed low-complexity regions of the assembly using DustMasker v.1.0.0 and identified candidate ORFs using TransDecoder v.5.7 (https://github.com/TransDecoder/TransDecoder/).^31^

### Assembly Completeness and Enzyme Annotation

The completeness of the assembly was investigated using Benchmarking Universal Single-Copy Orthologues (BUSCO) v2.0 with embryophyte (odb10) as the reference.^32^ A completeness investigation was also performed for the assembled genome and transcriptome data. Genetic data in which more than 42.5% of BUSCO sequences were missing were removed,^33^ resulting in a refined dataset comprising 1,434 species (appendix 1). Furthermore, we performed a focused search of specific enzyme families using BLAST v.2.12.0. We also performed a BLAST search for well-known specific groups of enzymes responsible for the biosynthesis of shikonin including PHB geranyltransferase (PGT), shikonin O-acyltransferase (SAT), geranylhydroquinone hydroxylase (GHQH), and deoxyshikonin hydroxylase (DSH), and benzylisoquinoline alkaloids including norcoclaurine 6-O-methyltransferase (6-OMT), coclaurine N -methyltransferase (CNMT), and 3′-hydroxy-N-methylcoclaurine 4′′-O-methyltransferase (4′-OMT). The specified identity threshold for each enzyme is shown in appendix 3 pp 4–21, with a bit score threshold set at 50.

### Spatial Pattern of Species Richness

We used the dataset of the occurrence of woody angiosperms compiled by ^24^. Focusing on species with more than ten occurrence records, we estimated the species distribution of our list of species with genome or transcriptome data passing completeness assessment. We specifically predicted species distributions at a scale of 1 degree per grid cell. A species distribution model was developed using Maxent version 3.4.1 with environmental variables, which included climatic (bio1–bio19), soil, geological, topographical, and geographical conditions, as the predictor variables (appendix 3 pp 22– 23).^34^ The relative suitability was computed within the range of countries where each species has a native distribution; the tree distribution checklist of each country was used to exclude nonnative distribution occurrence records. The Maxent output (maximum training sensitivity plus specificity Cloglog threshold) is a continuous map with shading intensities ranging from 0 to 1, allowing fine distinctions to be made between the modelled suitability of different areas. We used the threshold criteria of the sensitivity-specificity sum maximizer (MST) to generate a binary (presence/absence) prediction map for each species and confirmed the accuracy of each model using the area under the receiver operating characteristic curve (appendix 2).^35^ The relative suitability below the MST threshold value was converted to zero. We superimposed the distributions of the species with genome or transcriptome data and summed the number of species per grid cell. Finally, to assess the completeness of the geospatial coverage of sequenced species, we compared our diversity map to a map based solely on species occurrence data, as previously generated by ^24^ using a dataset comprising 82,974 species occurrences.

### Estimation of Natural Products Richness Based on Selected Marker

We estimated the richness of NPs by interpolating the average of 166 key enzymes within the grids. This was done by summing the number of enzyme markers identified from BLAST matches for all species present within the grid cell and dividing it by the number of species (Figs. 1 and 2B). Despite this cumulative average, we also mapped the distribution patterns of individual markers across the 166 enzymes (Fig. 3). We ranked the heterogeneity of each enzyme across different geographical regions by calculating the ratio of the maximum to the mean of the average enzyme count across all grids. Concurrently, the prevalence of the enzymes was estimated by the product of the total number of enzymes and the species count across all grid regions. Then, we created a scatterplot based on heterogeneity and enzyme prevalence to pinpoint any key enzymes that show an uneven and localized distribution across diverse geographical regions (Fig. 3F). In the biosynthetic pathways of shikonin derivatives and benzylisoquinoline alkaloids, distinct sets of enzymes are identified as pivotal to their synthesis. For shikonin, these include PGT, SAT, GHQH, and DSH, while for benzylisoquinoline alkaloids, the key enzymes are 6-OMT, CNMT, and 4′-OMT. To quantify the average enzymatic activity associated with these pathways, we performed an interpolation of data exclusively for species that possess the complete array of these specified enzymes. The average for each enzyme group was computed from the data of species containing the full enzyme repertoire. Subsequently, these calculated averages were plotted on a grid to facilitate the visualization and comparative analysis.

**Fig. 2.**
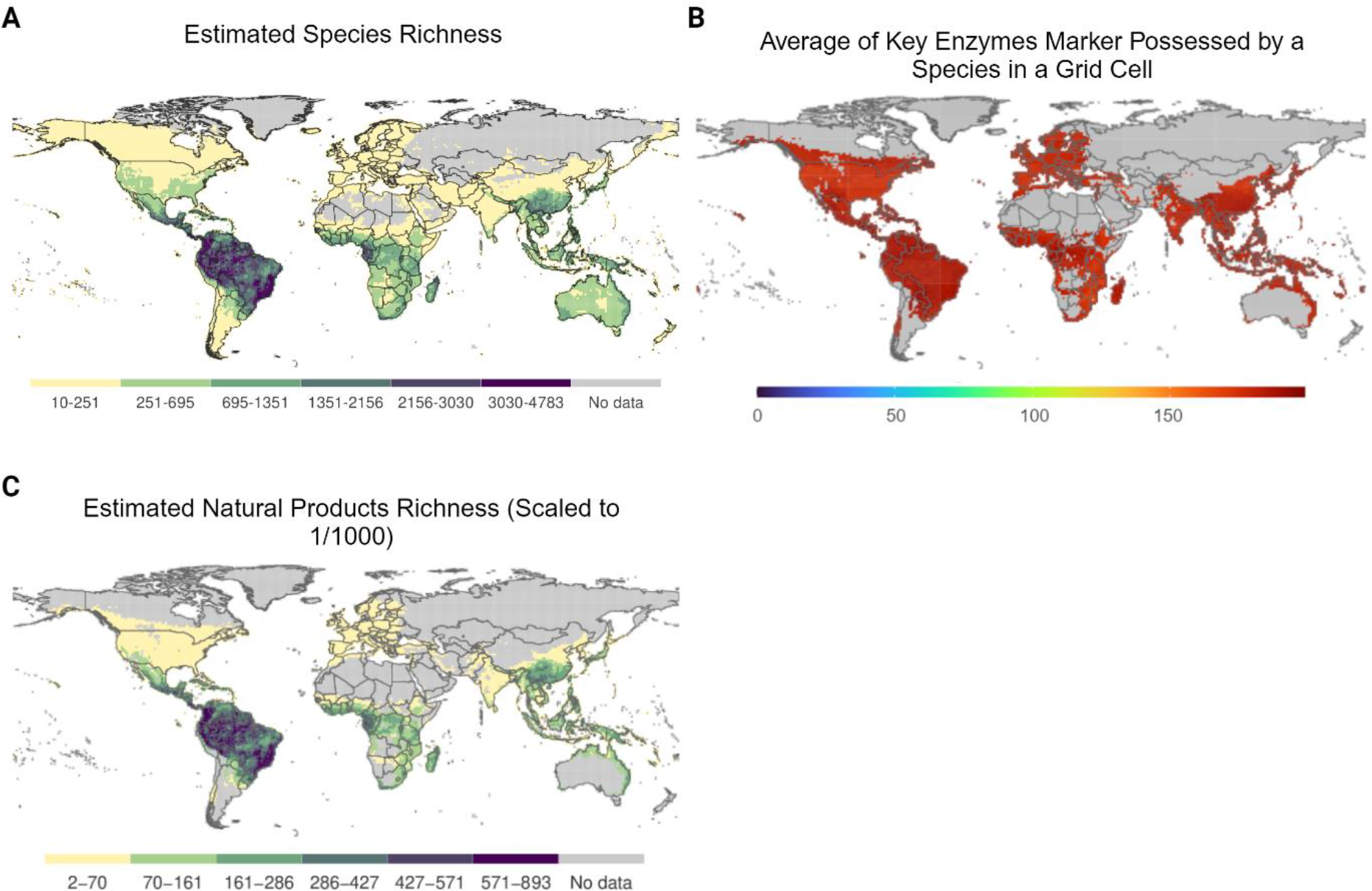
Global pattern of woody angiosperm species richness and genetic diversity. (A) The predicted species richness of woody angiosperm from 82,974 species. (B) The average of key enzyme markers for evaluating natural product diversity. (C) The predicted richness of natural products is derived from the product of average enzyme marker values and species richness.

**Fig. 3.**
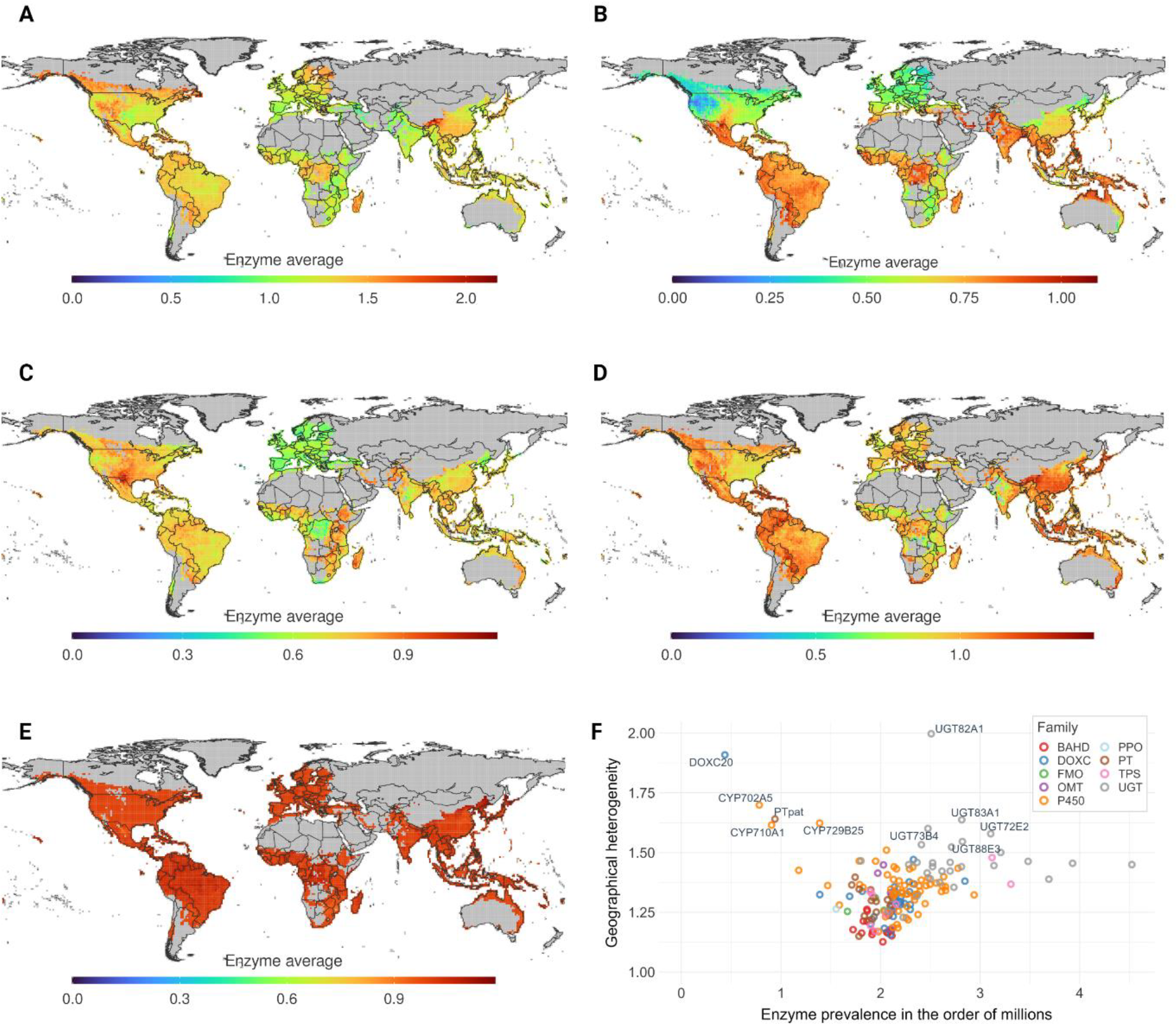
Global pattern of selected enzymes, including (A) UGT83A1, (B) CYP710A1, (C) PTpat, (D) CYP719A23, and (E) TcBAPT, and geographical heterogeneity and prevalence of enzyme markers (F).

## Results

The results show a general distribution of biodiversity; biodiversity hotspots are located near the equator, particularly in the South American Amazon, East Africa, and Southeast Asia (Fig. 2A). We employed the strategy of using the average of selected 166 key enzymes in a grid as a proxy to estimate NP hotspots or richness. Our result showed the presence of markers is evenly distributed globally (Fig. 2B), which, when paired with species richness, suggests that potential NP hotspots coincide with areas of high biodiversity (Fig. 2C). Furthermore, we conducted a detailed analysis of individual markers to identify the distribution patterns of each key enzymes. We found that several enzyme markers, such as UDP glycosyltransferase 82A1 (UGT82A1), 2-oxoglutarate-dependent dioxygenase-like protein (DOXC20), CYP702A5, 8-prenyl-1,3,6,7-tetrahydroxyxanthone 8-prenyltransferase (PTpat), and UGT83A1 (Fig. 3F), exhibit high geographical heterogeneity scores, indicating a disparate distribution contrary to Fig 2C. The remarkable geographical heterogeneity observed in the distribution of specific enzymes may hold profound implications for the spatial patterns of the corresponding NPs they synthesize. While the functional roles of UGT82A1, CYP702A5, and DOXC20 still warrant investigation, the biological attributes of UGT83A1, CYP710A1, and PTpat (Figs. 3A, B, and C) have been explored in previous studies. UGT83A1 has been implicated in the biosynthesis of sinapoyl anthocyanin, a compound with potent antioxidant properties.^36,37^ PTpat is involved in the biosynthesis of patulone, a compound noted for its anti-inflammatory properties.^38^ In addition, CYP710A1 is involved in the synthesis of stigmasterol from beta-sitosterol via C22 desaturation. Stigmasterol is suggested to possess anticancer and anti-inflammatory properties.^39,40^ Conversely, we can also find several enzymes that are linked to potential lead compound biosynthesis and show even geographical distribution. These include, for example, CYP719A23 and Taxus cuspidata Baccatin III:3-amino-3-phenylpropanoyltransferase (TcBAPT) (Figs. 3D and E). The CYP719 family has a distinct ability to produce a methylenedioxy ring, a characteristic chemical structure in etoposide, which is used to treat various cancers.^41,42^ The methylenedioxy rings catalyzed by the CYP719 family are also used in other bioactive compounds, including piperine and berberine.^43,44^ Meanwhile, TcBAPT is involved in paclitaxel biosynthesis, a widely used chemotherapy medication in cancer treatment.^45^

In light of the broad trends revealed by our initial analysis, we refined our methodology to focus on specific subsets of enzymes involved in the biosynthetic pathways of specific NPs. Our analyses showed that employing a targeted strategy allows for a more refined comprehension of NP distribution. For instance, the distribution of shikonin derivatives, potent natural compounds, presents an intriguing geographical constraint, with shikonin biosynthesis being concentrated predominantly in specific regions (Fig. 4A). These regions encompass Japan, South Korea, Taiwan, northern Vietnam, and areas along the Himalayan range. This pattern may reflect distinct evolutionary adaptations or specific ecological niches inhabited by plant species in these regions that promote the biosynthesis of shikonin derivatives. The confinement of shikonin biosynthesis to these locales emphasizes the crucial role of biodiversity conservation in these specific regions. In contrast, the distribution of benzylisoquinoline alkaloids do not exhibit such pronounced heterogeneity (Fig. 4B). Instead, its biosynthesis is more evenly distributed across different regions globally. This broad geographic distribution may be indicative of a more generalized ecological adaptation or a wider range of habitats conducive to the biosynthesis of benzylisoquinoline alkaloids. The ubiquity of benzylisoquinoline alkaloids biosynthesis and the constrained distribution of shikonin derivatives underscore the complexity and diversity of NP biosynthesis pathways and how they can be differentially influenced by the environmental, ecological, and evolutionary contexts of different regions. This finding further illustrates the importance of considering both the species and enzyme richness in a region while assessing the potential for medicinal resources in nature.

**Fig. 4.**
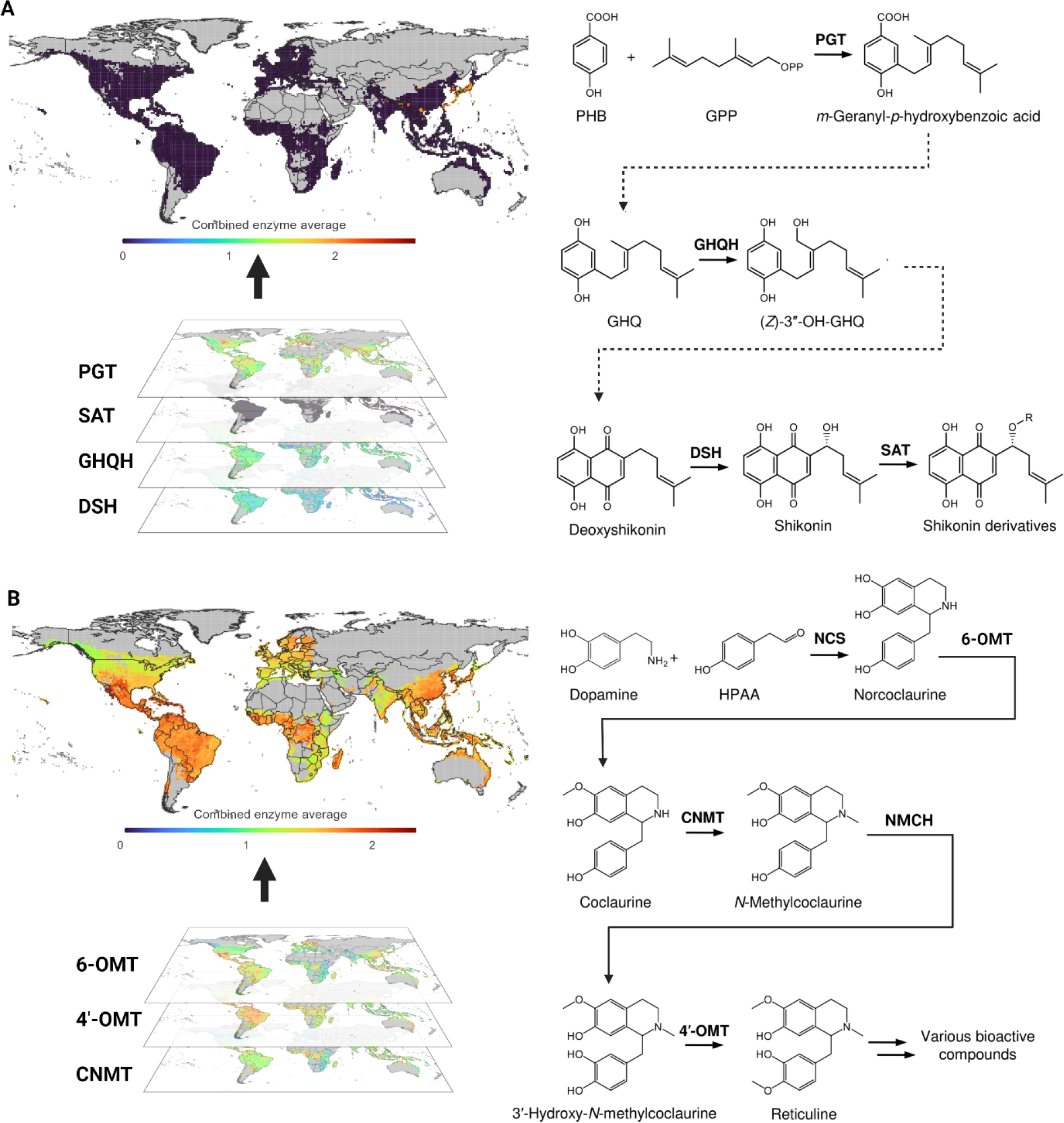
Global pattern of enzymes associated with shikonin and benzylisoquinoline biosynthetic pathways. In the biosynthetic pathway, dashed arrows indicate the reaction steps for which enzymes have not yet been identified. PHB, p-hydroxybenzoic acid; GPP, geranyl diphosphate; GHQ, geranylhydroquinone; (Z)-3^*′′*^-OH-GHQ, (Z)-3^*′′*^-hydroxy-geranylhydroquinone; PGT, PHB geranyltransferase; GHQH, geranylhydro-quinone hydroxylase; DSH, deoxyshikonin hydroxylase; SAT, shikonin O-acyltransferase; HPPA, 4-hydroxyphenylacetaldehyde; NCS, norcoclaurine synthase; 6-OMT, norcoclaurine 6-O-methyltransferase; CNMT, coclaurine N-methyltransferase; N-methylcoclaurine 3^*′*^-hydroxylase; 4^*′*^-OMT, 3-hydroxy-N-methylcoclaurine 4^*′*^-O-methyltransferase. (A) Geographic distribution of enzymes linked to shikonin biosynthesis, primarily concentrated in regions such as Japan, South Korea, Taiwan, Northern Vietnam, and areas around the Himalayas, suggesting region-specific ecological or evolutionary influences. The map is developed by overlaying the distributions of PGT, SAT, GHQH, and DSH. (B) Widespread distribution of enzymes associated with benzylisoquinoline biosynthesis. The map is developed by overlaying the distributions of 6-OMT, CNMT, and 4^*′*^-OMT.

## Discussion

The global environmental crisis in modern times threatens every aspect of biodiversity, ranging from individual genes to entire ecosystems.^46^ With limited conservation resources, a thorough biodiversity evaluation becomes crucial for directing targeted conservation efforts effectively. Despite the critical role of genomic in all biological domains,^47^ its integration into biodiversity assessments for conservation purposes remains insufficiently explored. Our study addresses this gap by integrating genomic and transcriptomic data with global distribution patterns to pinpoint hotspots for the conservation of NPs, which hold the medicinal resources. We discovered that key enzyme markers, which are indicative of NPs, exhibit a uniform global distribution (Fig. 2B). This uniformity, combined with data on species richness, leads us to propose that areas likely to be NP hotspots align with regions of high biodiversity (Fig. 2C). Our findings resonate with the principles outlined by ^48^ regarding marine natural products, extending the observed association between chemical diversity and biodiversity from marine biogeographical hotspots to terrestrial ecosystems, particularly within woody angiosperm hotspots. This insight suggests that protecting areas of high biodiversity should intrinsically conserve a broad array of NPs.

However, further analysis of the distribution of 166 enzyme markers revealed variable patterns. While the majority of markers showed a widespread distribution, mirroring the aggregate trends mentioned above, a subset exhibited a more localized presence, indicating that certain regions may serve as unique repositories for specific NPs (Fig. 3). This was exemplified when assessing group of enzymes involved in the biosynthesis of shikonin derivatives which also displayed distinctive, region-specific patterns (Fig. 4A). This variability in enzyme marker distribution underlines the complexity of NP biosynthesis and its ecological underpinnings. The widespread distribution of the majority of markers hints at a fundamental role these enzymes and their corresponding NPs play in broad ecological processes, possibly contributing to the adaptability and resilience of a variety of species across different environments. On the other hand, the localized distribution of certain markers points to a specialized ecological niche or adaptive strategy, wherein these specific NPs may confer selective advantages to woody angiosperm in particular habitats. Therefore, to effectively tailor conservation strategies, an understanding of the distinct distribution patterns of NP-related enzyme markers is essential.

The distribution of numerous NPs, which are highly localized, links them to potential risks from regional environmental changes. These risks can undermine their multifaceted roles. For instance, we found that the combined spatial patterns of enzymes associated with the biosynthesis of shikonin derivatives are predominantly localized to East Asian regions and areas along the Himalayan range. These compounds are known for their therapeutic potential in treating cancer and inflammation, as well as their traditional role in wound healing.^49^ Remarkably, shikonin derivatives, which accumulate exclusively within the roots of plants, also demonstrate potent antibacterial activity against soil-borne microorganisms.^50^ This particular characteristic, functioning as a chemical barrier protecting root tissues from biological stressors, may reflect unique regional evolutionary adaptations or ecological niches favouring shikonin derivative biosynthesis, a hypothesis warranting further investigation. The presence of these versatile NPs in specific regions underscores the importance of targeted conservation strategies.

The growing popularity of certain plants for use as herbal medicines or as sources of bioactive compounds for pharmaceutical drugs often leads to a surge in global market demand.^51^ This increase can result in overharvesting from natural populations, putting the species at risk of extinction.^51,52^ It is estimated that between 60% and 80% of medicinal plants traded worldwide are gathered from the wild, representing a trade value that surpasses $3 billion.^51^ Additionally, the distribution of medicinal plants throughout the world is unequal, with China and India leading the way in terms of utilization, with 11,146 and 7,500 species, respectively.^53^ Understanding the spatial patterns of NPs may open avenues for influencing market dynamics. If overharvesting is a concern in regions that need conservation, recommendations for alternative medicinal plants with similar NP profiles can be made. These approaches have the potential to alleviate pressure on endangered or overharvested regions by shifting the demand in the market towards more sustainable sources.

While our study provides significant insights into the spatial patterns of enzymes relevant to NP with medicinal properties, several limitations should be acknowledged. Our study primarily focused on woody angiosperm, which leaves out a significant portion of plant biodiversity, such as herbaceous plants, grasses, and aquatic plants. This limited scope may consequently exclude potential drug candidates found within these groups. Additionally, the quality and completeness of the genetic and distributional data can affect the accuracy of our results (appendix 3 p 24). Incomplete or biased data could potentially lead to the misinterpretation or oversight of significant patterns. While we found significant geospatial heterogeneity in numerous enzymes, these enzymes alone do not directly translate into NP. Finally, our approach might not capture the complexities of plant biochemical pathways. Future research that addresses these gaps will broaden the application of our findings to the real world.

## Contributors

YK, YKakei, KT and KK contributed to the conceptualisation of the study and the supervision. MF and KK contributed to the writing – original draft. All authors contributed to the methodology, investigation and writing – review & editing. MF and KT contributed to the visualization. KK contributed to the funding acquisition and project administration.

## Declaration of interests

We declare no competing interests.

## Data sharing

All data used in our analysis are available in public databases and appendixes.

## Acknowledgments

This work is supported by Japan Society for the Promotion of Science through the Grant-in-Aid for Scientific Research (A) 20H00651.

## Notes

### Competing Interest Statement

The authors have declared no competing interest.

### Summary of Updates

Figure illustrating the design of the study added (Figure 2). Figure 2 and 3 combined into one figure (figure 3). Section on abstract and discussion updated to clarify the aim of the study.

## References

1. S. Bernardini, A. Tiezzi, V. L. Masci, E. Ovidi, Natural products for human health: An historical overview of the drug discovery approaches. Natural Product Research 32, 1926–1950 (2018). doi:10.1080/14786419.2017.1356838

2. G. M. Cragg, D. J. Newman, Natural products: A continuing source of novel drug leads. Biochimica et Biophysica Acta (BBA) - General Subjects 1830, 3670–3695 (2013). doi: 10.1016/j.bbagen.2013.02.008

3. D. J. Newman, G. M. Cragg, Natural products as sources of new drugs over the nearly four decades from 01/1981 to 09/2019. Journal of Natural Products 83, 770–803 (2020). doi:10.1021/acs.jnatprod.9b01285

4. G. Karageorgis, D. J. Foley, L. Laraia, H. Waldmann, Principle and design of pseudo-natural products. Nature Chemistry 12, 227–235 (2020). doi:10.1038/s41557-0190411-x

5. X. Zhang, S. Li, Expansion of chemical space for natural products by uncommon P450 reactions. Natural Product Reports 34, 1061–1089 (2017). doi:10.1039/C7NP00028F

6. S. Jennewein, R. M. Long, R. M. Williams, R. Croteau, Cytochrome P450 taxadiene 5alpha-hydroxylase, a mechanistically unusual monooxygenase catalyzing the first oxygenation step of taxol biosynthesis. Chemistry & Biology 11, 379–387 (2004). doi:10.1016/j.chembiol.2004.02.022

7. K. H. Teoh, D. R. Polichuk, D. W. Reed, G. Nowak, P. S. Covello, Artemisia annua L. (Asteraceae) trichome-specific cDNAs reveal CYP71AV1, a cytochrome P450 with a key role in the biosynthesis of the antimalarial sesquiterpene lactone artemisinin. FEBS Letters 580, 1411–1416 (2006). doi:10.1016/j.febslet.2006.01.065

8. J. A. Chemler, M. A. Koffas, Metabolic engineering for plant natural product biosynthesis in microbes. Current Opinion in Biotechnology 19, 597–605 (2008). doi:10.1016/j.copbio.2008.10.011

9. B. J. Cardinale, J. E. Duffy, A. Gonzalez, D. U. Hooper, C. Perrings, P. Venail, A. Narwani, G. M. Mace, D. Tilman, D. A. Wardle, A. P. Kinzig, G. C. Daily, M. Loreau, J. B. Grace, A. Larigauderie, D. S. Srivastava, S. Naeem, Biodiversity loss and its impact on humanity. Nature 486, 59–67 (2012). doi:10.1038/nature11148

10. IUCN, The IUCN red list of threatened species. Version 2022-2. https://www.iucnredlist.: Accessed 2022-10-30

11. M. C. Hansen, P. V. Potapov, R. Moore, M. Hancher, S. A. Turubanova, A. Tyukavina, D. Thau, S. V. Stehman, S. J. Goetz, T. R. Loveland, A. Kommareddy, A. Egorov, L. Chini, C. O. Justice, J. R. G. Townshend, High-resolution global maps of 21st-century forest cover change. Science 342, 850–853 (2013). doi:10.1126/science.1244693

12. P. G. Curtis, C. M. Slay, N. L. Harris, A. Tyukavina, M. C. Hansen, Classifying drivers of global forest loss. Science 361, 1108–1111 (2018). doi:10.1126/science.aau3445

13. K. J. Gaston, Global patterns in biodiversity. Nature 405, 220–227 (2000). doi:10.1038/35012228

14. M. R. Willig, D. M. Kaufman, R. D. Stevens, Latitudinal gradients of biodiversity: Pattern, process, scale, and synthesis. Annual Review of Ecology, Evolution, and Systematics 34, 273–309 (2003). doi:10.1146/annurev.ecolsys.34.012103.144032

15. R. L. Pressey, R. M. Cowling, M. Rouget, Formulating conservation targets for biodiversity pattern and process in the cape floristic region, south africa. Biological Conservation 112, 99–127 (2003). doi:10.1016/S0006-3207(02)00424-X.

16. X. Irigoien, J. Huisman, R. P. Harris, Global biodiversity patterns of marine phytoplankton and zooplankton. Nature 429, 863–867 (2004). doi:10.1038/nature02593

17. L. Mair et al., A metric for spatially explicit contributions to science-based species targets. Nature Ecology & Evolution 5, 836–844 (2021). doi:10.1038/s41559-021-01432-0

18. A. Miraldo et al., An anthropocene map of genetic diversity. Science 353, 1532–1535 (2016). doi:10.1126/science.aaf4381

19. H. De Kort et al., Life history, climate and biogeography interactively affect worldwide genetic diversity of plant and animal populations. Nature Communications 12, 516 (2021). doi:10.1038/s41467-021-20958-2

20. S. Theodoridis et al., Evolutionary history and past climate change shape the distribution of genetic diversity in terrestrial mammals. Nature Communications 11, 2557 (2020). doi:10.1038/s41467-020-16449-5

21. N. R. Owen et al., Global conservation of phylogenetic diversity captures more than just functional diversity. Nature Communications 10, 859 (2019). doi:10.1038/s41467-019-08600-8

22. S. Yadav, A. Sharma, G.A. Nayik, R. Cooper, G. Bhardwaj, H.S. Sohal, V. Mutreja, R. Kaur, F.O. Areche, M. AlOudat, A.M. Shaikh, B. Kovács, A.E. Mohamed Ahmed, Review of shikonin and derivatives: Isolation, chemistry, biosynthesis, pharmacology and toxicology. Frontiers in Pharmacology 13, (2022). doi:10.3389/fphar.2022.905755

23. A. Vadhel, S. Bashir, A.H. Mir, M. Girdhar, D. Kumar, A. Kumar, A. Mohan, T. Malik, A. Mohan, Opium alkaloids, biosynthesis, pharmacology and association with cancer occurrence. Open Biology 13, 220355 (2023). doi:10.1098/rsob.220355

24. B. Kusumoto, A. Chao, W. L. Eiserhardt, J .-C. Svenning, T. Shiono, Y. Kubota, Occurrence-based diversity estimation reveals macroecological and conservation knowledge gaps for global woody plants. Science Advances 9(40), 9719 (2023). DOI:10.1126/sciadv.adh9719

25. N. Matasci, L. -H. Hung, Z. Yan, E. J. Carpenter, N. J. Wickett, S. Mirarab, N. Nguyen, T. Warnow, S. Ayyampalayam, M. Barker, J. G. Burleigh, M. A. Gitzendanner, E. Wafula, J. P. Der, C. W. dePamphilis, B. Roure, H. Philippe, B. R. Ruhfel, N. W. Miles, S. W. Graham, S. Mathews, B. Surek, M. Melkonian, D. E. Soltis, P. S. Soltis, C. Rothfels, L. Pokorny, J. A. Shaw, L. DeGironimo, D. W. Stevenson, J. C. Villarreal, T. Chen, T. M. Kutchan, M. Rolf, R. S. Baucom, M. K. Deyholos, R. Samudrala, Z. Tian, X. Wu, X. Sun, Y. Zhang, J. Wang, J. Leebens-Mack, G. K. -S. Wong, Data access: For the 1,000 Plants (1KP) project. GigaScience 3, 2047–2173 (2014). doi:10.1186/2047-217X-3-17

26. S. Chen, Y. Zhou, Y. Chen, J. Gu, Fastp: An ultra-fast all-in-one FASTQ preprocessor. Bioinformatics 34, 884–890 (2018). doi:10.1093/bioinformatics/bty560

27. P. Ewels, M. Magnusson, S. Lundin, M. Käller, MultiQC: Summarize analysis results for multiple tools and samples in a single report. Bioinformatics 32, 3047–3048 (2016). doi:10.1093/bioinformatics/btw354

28. E. Kopylova, L. Noé, H. Touzet, SortMeRNA: Fast and accurate filtering of ribosomal RNAs in metatranscriptomic data. Bioinformatics 28, 3211–3217 (2012). doi:10.1093/bioinformatics/bts611

29. Y. Xie, G. Wu, J. Tang, R. Luo, J. Patterson, S. Liu, W. Huang, G. He, S. Gu, S. Li, X. Zhou, T. -W. Lam, Y. Li, X. Xu, G. K. -S. Wong, J. Wang, SOAPdenovo-Trans: De novo transcriptome assembly with short RNA-Seq reads. Bioinformatics 30, 1660–1666 (2014). doi:10.1093/bioinformatics/btu077

30. P. Di Tommaso, M. Chatzou, E. W. Floden, P. P. Barja, E. Palumbo, C. Notredame, Nextflow: Enables reproducible computational workflows. Nature Biotechnology 35, 316–319 (2017). doi:10.1038/nbt.3820

31. A. Morgulis, E. M. Gertz, A. A. Schäffer, R. Agarwala, A fast and symmetric DUST implementation: To mask low-complexity DNA sequences. Journal of Computational Biology 13, 1028–1040 (2006). doi:10.1089/cmb.2006.13.1028

32. F.A. Simão, R.M. Waterhouse, P. Ioannidis, E.V. Kriventseva, E.M. Zdobnov, BUSCO: Assessing genome assembly and annotation completeness with single-copy orthologs. Bioinformatics 31, 3210–3212 (2015). doi:10.1093/bioinformatics/btv351

33. J.H. Leebens-Mack, et al., One thousand plant transcriptomes and the phylogenomics of green plants. Nature 574, 679–685 (2019). doi:10.1038/s41586-019-1693-2

34. S. J. Phillips, R. P. Anderson, R. E. Schapire, Maximum entropy modeling of species geographic distributions. Ecological Modelling 190, 231–259 (2006). doi:10.1016/j.ecolmodel.2005.03.026

35. A. Jiménez-Valverde, J. M. Lobo, Threshold criteria for conversion of probability of species presence to either–or presence–absence. Acta Oecologica 31, 361–369 (2007). doi:10.1016/j.actao.2007.02.001

36. Z. Ren et al., Network analysis of transcriptome and LC-MS reveals a possible biosynthesis pathway of anthocyanins in Dendrobium officinale. BioMed Research International 2020, 6512895 (2020). doi:10.1155/2020/6512895

37. Z. Xiao, L. Fang, Y. Niu, H. Yu, Effect of cultivar and variety on phenolic compounds and antioxidant activity of cherry wine. Food Chemistry 186, 69–73 (2015). doi:10.1016/j.foodchem.2015.01.050

38. M. Nagia et al., Sequential regiospecific gem-diprenylation of tetrahydroxyxanthone by prenyltransferases from Hypericum sp. New Phytologist 222, 318–334 (2019). doi:10.1111/nph.15611

39. T. Griebel, J. Zeier, A role for beta-sitosterol to stigmasterol conversion in plant– pathogen interactions. The Plant Journal 63, 254–268 (2010). doi:10.1111/j.1365-313X.2010.04235.x

40. S. Bakrim, N. Benkhaira, I. Bourais, T. Benali, L.-H. Lee, N. El Omari, R.A. Sheikh, K.W. Goh, L.C. Ming, A. Bouyahya, Health benefits and pharmacological properties of stigmasterol. Antioxidants 11(10) (2022). 10.3390/antiox11101912

41. J. V. Marques, K. W. Kim, C. Lee, M. A. Costa, G. D. May, J. A. Crow, L. B. Davin, N. G. Lewis, Next generation sequencing in predicting gene function in podophyllotoxin biosynthesis. Journal of Biological Chemistry 288, 466–479 (2013). doi:10.1074/jbc.M112.400689

42. W. Lau, E. S. Sattely, Six enzymes from mayapple that complete the biosynthetic pathway to the etoposide aglycone. Science 349, 1224–1228 (2015). doi:10.1126/science.aac7202

43. A. Schnabel, F. Cotinguiba, B. Athmer, T. Vogt, Piper nigrum CYP719A37 catalyzes the decisive methylenedioxy bridge formation in piperine biosynthesis. Plants 10, 128 (2021). doi:10.3390/plants10010128

44. N. Ikezawa, M. Tanaka, M. Nagayoshi, R. Shinkyo, T. Sakaki, K. Inouye, F. Sato, Molecular cloning and characterization of CYP719, a methylenedioxy bridge-forming enzyme that belongs to a novel P450 family, from cultured Coptis japonica cells. Journal of Biological Chemistry 278, 38557–38565 (2003). doi:10.1074/jbc.M302470200

45. K. Walker, R. Long, R. Croteau, The final acylation step in taxol biosynthesis: cloning of the taxoid C13-side-chain N-benzoyltransferase from Taxus. Proceedings of the National Academy of Sciences 99, 9166–9171 (2002). doi:10.1073/pnas.082115799

46. W. Steffen, K. Richardson, J. Rockström, S.E. Cornell, I. Fetzer, E.M. Bennett, R. Biggs, S.R. Carpenter, W. de Vries, C.A. de Wit, C. Folke, D. Gerten, J. Heinke, G.M. Mace, L.M. Persson, V. Ramanathan, B. Reyers, S. Sörlin, Planetary boundaries: Guiding human development on a changing planet. Science 347, 1259855 (2015). doi:10.1126/science.1259855

47. K. Theissinger et al., How genomics can help biodiversity conservation. Trends in Genetics 39, 545–559 (2023). doi:10.1016/j.tig.2023.01.005

48. M.E. Hay, Marine chemical ecology: what’s known and what’s next? Journal of Experimental Marine Biology and Ecology 200(1), 103–134 (1996). 10.1016/S0022-0981(96)02659-7

49. C. Guo, J. He, X. Song, L. Tan, M. Wang, P. Jiang, Y. Li, Z. Cao, C. Peng, Pharmacological properties and derivatives of shikonin—a review in recent years. Pharmacological Research 149, 104463 (2019). doi:10.1016/j.phrs.2019.104463

50. L. A. Brigham, P. J. Michaels, H. E. Flores, Cell-Specific Production and Antimicrobial Activity of Naphthoquinones in Roots of Lithospermum erythrorhizon. Plant Physiology 119, 417–428 (1999). doi:10.1104/pp.119.2.417

51. S. Theodoridis, E. G. Drakou, T. Hickler, M. Thines, D. Nogues-Bravo, Evaluating natural medicinal resources and their exposure to global change. The Lancet Planetary Health 7, 155–163 (2023). doi:10.1016/S2542-5196(22)00317-5

52. M. -J. Howes, C. Quave, J. Collemare, E. Tatsis, D. Twilley, E. Lulekal, A. Farlow, L. al, E. Bell, Molecules from nature: Reconciling biodiversity conservation and global healthcare imperatives for sustainable use of medicinal plants and fungi. Plants People Planet 2, 463–481 (2020). doi:10.1002/ppp3.10138

53. S. -L. Chen, H. Yu, H. -M. Luo, Q. Wu, C. -F. Li, A. Steinmetz, Conservation and sustainable use of medicinal plants: problems, progress, and prospects. Chinese Medicine 11, 37 (2016). doi:10.1186/s13020-016-0108-7

