## Appendix 3 for "Mapping Natural Product Biosynthetic Hotspots: Prioritizing Conservation for Medicinal Resources"

*Supporting Appendix 3*

**Mapping Natural Product Biosynthetic Hotspots: Prioritizing Conservation for Medicinal Resources**

Muhamad Fahmi^1^*, Kojiro Takanashi^2,3^, Yusuke Kakei^4^, Yasuhiro Kubota^5,6^, Keiichiro Kanemoto^1,7,8^*

^1^Research Institute for Humanity and Nature, 457-4 Motoyama, Kamigamo, Kita-ku, Kyoto, 603-8047, Kyoto, Japan.

^2^Department of Science, Graduate School of Science and Technology, Shinshu University, 3-1-1 Asahi, Matsumoto, 390-8621, Nagano, Japan.

^3^Department of Biology, Faculty of Science, Shinshu University, 3-1-1 Asahi, Matsumoto, 390-8621, Nagano, Japan.

^4^The Institute of Vegetable and Floriculture Science, National Agriculture and Food Research Organization, 3-1-1 Kannondai, Tsukuba, 244-0813, Ibaraki, Japan.

^5^Faculty of Science, University of the Ryukyus, 1, Sembaru, Nishihara, 903-0213, Okinawa, Japan.

^6^Think Nature Inc., Nishihara, 903-0213, Okinawa, Japan.

^7^Graduate School of Environmental Studies, Tohoku University, Aoba, 468-1, Aramaki, Aoba-ku, Sendai, 980-8572, Miyagi, Japan. ^8^Research Center for Social Systems, Shinshu University, 3-1-1 Asahi, Matsumoto, 390-8621, Nagano, Japan.

Contributing authors:

### Table of contents

**Table S1**. Comprehensive enzyme information with emphasis on functions, sequences, and blast identity thresholds.	S4

**Table S2**. Comprehensive enzyme information associated to shikonin and benzylisoquinoline biosynthesis with emphasis on functions, sequences, and blast identity thresholds	S21

**Table S3**. Environmental data used for distribution prediction by MaxEnt.	S22

**Figure S1.** The completeness of the genetic and distributional data	S24

**Figure S2.** Global pattern of (A) GmIF7MaT, (B) Pf5MaT, (C) At3AT1, (D) ZmGlossy2, (E) CER2, (F), CrDAT, (G) FaSAAT, (H) PsSalAT, (I) PhCFAT, and and (J) HvACT	S25

**Figure S3.** Global pattern of (A) HHT1, (B) TcBAPT, (C) LaHMT/HLT, (D) TpHCT2, (E) AtHCT, (F) DOXC3, (G) DOXC7, (H) DOXC12, (I) DOXC13, and (J) DOXC14	S26

**Figure S4.** Global pattern of (A) DOXC15, (B) DOXC17, (C) DOXC19, (D) DOXC20, (E) DOXC21, (F) DOXC23, (G) DOXC24, (H) DOXC27, (I) DOXC28, and (J) DOXC30	S27

**Figure S5.** Global pattern of (A) DOXC31, (B) DOXC31, (C) DOXC37, (D) DOXC38, (E) DOXC45, (F) DOXC46, (G) DOXC47, (H) DOXC47, (I) DOXC52, and (J) DOXC53	S28

**Figure S6.** Global pattern of (A) DOXC54, (B) DOXC55, (C) GSXL7, (D) FMO2, (E) YUC7, (F) CAMT4, (G) BSMT1, (H) FAMT, (I) TSM1, and (J) IGMT3	S29

**Figure S7.** Global pattern of (A) NAMT1, (B) OMT1, (C) CYP51G1, (D) CYP71D1V2, (E) CYP72A1, (F) CYP73A1, (G) CYP74D2, (H) CYP75B2, (I) CYP76M8, and (J) CYP77A2	S30

**Figure S8.** Global pattern of (A) CYP78A5, (B) CYP79A1, (C) CYP80G2, (D) CYP81E1, (E) CYP82Y1, (F) CYP83B1, (G) CYP84A4, (H) CYP85A3, (I) CYP86A1, and (J) CYP87D18	S31

**Figure S9.** Global pattern of (A) CYP88A3, (B) CYP89A2, (C) CYP90C1, (D) CYP91A2, (E) CYP92C5, (F) CYP93C2, (G) CYP94B3, (H) CYP95, (I) CYP96A15, and (J) CYP97C1	S32

**Figure S10.** Global pattern of (A) CYP98A8, (B) CYP99A3, (C) CYP701A8, (D) CYP702A5, (E) CYP703A2, (F) CYP704B1, (G) CYP705A5, (H) CYP706A3, (I) CYP707A2, and (J) CYP708A2	S33

**Figure S11.** Global pattern of (A) CYP709B3, (B) CYP710A1, (C) CYP711A1, (D) CYP712A1, (E) CYP714B1, (F) CYP715A1, (G) CYP716A47, (H) CYP718B1, (I) CYP719A23, and (J) CYP720B1	S34

**Figure S12.** Global pattern of (A) CYP721A1, (B) CYP722A1, (C) CYP724B1, (D) CYP725A1, (E) CYP726A13, (F) CYP727C1, (G) CYP728S3, (H) CYP729B25, (I) CYP733A1, and (J) CYP734A1	S35

**Figure S13.** Global pattern of (A) CYP735A2, (B) CYP736A117, (C) MdPPO, (D) COX10, (E) HPT_VTE2-1, (F) ClPT1, (G) N8DT-1, (H) ABC4, (I) CHLG, and (J) PPT1	S36

**Figure S14.** Global pattern of (A) HST, (B) PcPT, (C) AcPT1, (D) RdPT1, (E) HGGT, (F) PT1, (G) Ptpat, (H) FPT, (I) PGT-1, and (J) TPS26	S37

**Figure S15.** Global pattern of (A) TP20L, (B) AtTPS14, (C) TPS3, (D) TPS4, (E) TPS31, (F) TPS41, (G) smTPS4, (H) UGT71C1, (I) UGT72E2, and (J) UGT73B4	S38

**Figure S16.** Global pattern of (A) UGT74F2, (B) UGT75B1, (C) UGT76G1, (D) UGT77B2, (E) UGT78G1, (F) UGT79B1, (G) UGT80A2, (H) UGT81A1, (I) UGT82A1, and (J) UGT83A1	S39

**Figure S17.** Global pattern of (A) UGT84A1, (B) UGT85B1, (C) UGT86A2, (D) UGT87A2, (E) UGT88E3, (F) UGT89C1, (G) UGT90A1, (H) UGT91D2, (I) UGT92A1, and (J) UGT93B8	S40

**Figure S18.** Global pattern of (A) UGT94B1, (B) UGT95B1, (C) UGT97B3, (D) UGT98B4, (E) UGT99A6, and (F) UGT100	S41

**Section 1:** De novo assembly simulation to elucidate optimal k-mer	S42

**Table S1**. Comprehensive enzyme information with emphasis on functions, sequences, and blast identity thresholds

| Uniprot/NCBI ID | Enzyme | Family | Organism | Associated Compound | Potential use | PMID | Blast Identity threshold |
| --- | --- | --- | --- | --- | --- | --- | --- |
| Q9SAA9 | CYP51G1 | CYP | *Arabidopsis thaliana* | Ergosterol | Antifungal | 20547249; 8852337 | 40 |
| I1TEM1 | CYP71D1V2 | CYP | *Catharanthus roseus* | Vindoline; Anhydrovinblastine | Anti-tumor; Anti-diabetic | 24012527; 22115594; 25918424; 25850027; 22837051 | 40 |
| Q05047 | CYP72A1 | CYP | *Catharanthus roseus* | Secologanin; Ipecac alkaloids; Monoterpene indole alkaloids | Anti-cancer drug lead | 32745157; 33477682; 11135113; 26285573; 26285573 | 40 |
| Q04468 | CYP73A1 | CYP | *Helianthus tuberosus* | Phenylpropanoid | Plant defense | 8026495; 9398253; 14576280; 8026495; 9398253 | 40 |
| Q9AVQ1 | CYP74D2 | CYP | *Solanum tuberosum* | Colneleic acid; Colnelenic acid | Anti-fungal; Anti-microbial | 17085514; 11696374 | 40 |
| Q9SBQ9 | CYP75B2 | CYP | *Petunia hybrida* | Eriodictyol; Dihydroquercetin | Anti-inflammatory and antioxidant | 21299115; 24610848; 27250501 | 40 |
| Q6YTF1 | CYP76M8 | CYP | *Oryza sativa subsp. japonica* | Phytocassane; Oryzalexin | Anti-diabetes; Anti-cancer | 37010741 | 40 |
| P37124 | CYP77A2 | CYP | *Solanum melongena* | N/A | Anti-fungal | 32046212 | 40 |
| Q9LMX7 | CYP78A5 | CYP | *Arabidopsis thaliana* | Linalool oxides | Antidepressant; Antimicrobial; Anti-cancer | 34544345; 31379158; 34611674 | 40 |
| Q43135 | CYP79A1 | CYP | *Sorghum bicolor* | p-hydroxyphenylacetaldehyde oxime; Dhurrin | Natural pest control agent | 2250015; 7876084; 36279963 | 40 |
| A8CDR5 | CYP80G2 | CYP | *Coptis japonica* | Magnoflorine; Orientaline | Anti-inflammatory; Sedative and anxiolytic effects | 18230623; 23456265; 25840917 | 40 |
| P93147 | CYP81E1 | CYP | *Glycyrrhiza echinata* | 2'-hydroxyisoflavones, 2'-hydroxydaidzein, and 2'-hydroxyformononetin | Antimicrobial | 9790908 | 40 |
| I3PLR1 | CYP82Y1 | CYP | *Papaver somniferum* | Noscapine | Chemotherapeutic Agent; Antitussive; Anti-inflammatory effects; Antioxidant | 24324259; 27378283; 29610307; 34880922; 27237331 | 40 |
| O65782 | CYP83B1 | CYP | *Arabidopsis thaliana* | Indole glucosinolate | Plant disease resistance | 26361733; 28154137 | 40 |
| F4JW83 | CYP84A4 | CYP | *Arabidopsis thaliana* | Arabidopyrones; Caffealdehyde | N/A | 22923580 | 40 |
| Q50LE0 | CYP85A3 | CYP | *Solanum lycopersicum* | Brassinolide | Anti-cancer | 15710611; 28847728 | 40 |
| P48422 | CYP86A1 | CYP | *Arabidopsis thaliana* | Suberin | Drought-resistance gene; Plant defense | 18544608 | 40 |
| K7NBR2 | CYP87D18 | CYP | *Siraitia grosvenorii* | Mogrosides | Anti-leukemia; Anti-inflammatory | 26903528; 33296813; 26101699 | 40 |
| O23051 | CYP88A3 | CYP | *Arabidopsis thaliana* | Gibberellins | N/A | 11172076 | 40 |
| Q42602 | CYP89A2 | CYP | *Arabidopsis thaliana* | N/A | N/A | N/A | 40 |
| Q9M066 | CYP90C1 | CYP | *Arabidopsis thaliana* | Brassinolide | Anti-cancer | 17138693; 28847728 | 40 |
| A0A251UB05 | CYP91A2 | CYP | *Helianthus annuus* | N/A | N/A | N/A | 40 |
| A0A1D6HSP4 | CYP92C5 | CYP | *Zea mays* | Dimethylnonatriene; Trimethyltridecatetraene | Plant defense | 27662898; 30187155; 10482672 | 40 |
| Q9SXS3 | CYP93C2 | CYP | *Glycyrrhiza echinata* | Daidzein; Genistein | Anti-cancer; Anti-diabetes | 23870911; 10557230; 20116654 | 40 |
| Q9SMP5 | CYP94B3 | CYP | *Arabidopsis thaliana* | 12-hydroxy-JA-Ile | N/A | 21849397; 21576464 | 40 |
| Q8RWY7 | CYP95 | CYP | *Arabidopsis thaliana* | N/A | N/A | N/A | 40 |
| Q9FVS9 | CYP96A15 | CYP | *Arabidopsis thaliana* | N/A | N/A | 17905869 | 40 |
| Q6TBX7 | CYP97C1 | CYP | *Arabidopsis thaliana* | Lutein | Neuroprotective agent; Anti-inflammatory | 16492736; 34111488 | 40 |
| Q9CA61 | CYP98A8 | CYP | *Arabidopsis thaliana* | Spermidine | Protects against acute kidney injury | 19762055; 22303276; 36058905 | 40 |
| Q0JF01 | CYP99A3 | CYP | *Oryza sativa subsp. japonica* | Momilactone | Antifungal | 29996047 | 40 |
| Q0DBF4 | CYP701A8 | CYP | *Oryza sativa subsp. japonica* | Oryzalexin D and Oryzalexin E; Phytocassane | Anti-diabetes; Anti-cancer | 22247270; 23795884; 37010741 | 40 |
| A8MS53 | CYP702A5 | CYP | *Arabidopsis thaliana* | N/A | N/A | N/A | 40 |
| Q9LNJ4 | CYP703A2 | CYP | *Arabidopsis thaliana* | Sporopollenin | Microencapsulation; Drug delivery systems | 35929662; 35201750; 36244536 | 40 |
| Q9C788 | CYP704B1 | CYP | *Arabidopsis thaliana* | Sporopollenin | Microencapsulation; Drug delivery systems | 19700560; 35201750; 36244536 | 40 |
| Q9FI39 | CYP705A5 | CYP | *Arabidopsis thaliana* | Desaturated thalian-diol | N/A | 29170672 | 40 |
| Q9LU04 | CYP706A3 | CYP | *Arabidopsis thaliana* | Scutellein | Neuroprotective agent | 29386648 | 40 |
| K4CI52 | CYP707A2 | CYP | *Solanum lycopersicum* | N/A | N/A | N/A | 40 |
| Q8L7D5 | CYP708A2 | CYP | *Arabidopsis thaliana* | Thalian-diol | N/A | N/A | 40 |
| Q9T093 | CYP709B3 | CYP | *Arabidopsis thaliana* | N/A | N/A | 24164720 | 40 |
| O64697 | CYP710A1 | CYP | *Arabidopsis thaliana* | Stigmasterol | Anti-inflammatory; Anti-cancer; immunomodulatory; Neuroprotective agent | 20444228; 36290632 | 40 |
| B9DFU2 | CYP711A1 | CYP | *Arabidopsis thaliana* | Carlactonoic acid; Strigolactone | Anti-cancer; Antimicrobial agents | 25425668; 34361731 | 40 |
| O48532 | CYP712A1 | CYP | *Arabidopsis thaliana* | N/A | N/A | N/A | 40 |
| Q7XHW5 | CYP714B1 | CYP | *Oryza sativa subsp. japonica* | Gibberellins | N/A | N/A | 40 |
| F4KG63 | CYP715A1 | CYP | *Arabidopsis thaliana* | N/A | N/A | N/A | 40 |
| H2DH16 | CYP716A47 | CYP | *Panax ginseng* | Ginsenosides | Anti-cancer; Neuroprotective agent; Anti-inflammatory; Anti-diabetes | 22039120; 25981048; 36278171; 29854792; 32183094; 31835292 | 40 |
| A0A291FB27 | CYP718B1 | CYP | *Taxus wallichiana var. chinensis* | N/A | N/A | N/A | 40 |
| L7T8H2 | CYP719A23 | CYP | *Sinopodophyllum hexandrum* | Etoposide; Pluviatolide | Chemotherapeutic compound; Anti-cancer | 23161544; 23161544; 9893622; 23161544 | 40 |
| Q50EK6 | CYP720B1 | CYP | *Pinus taeda* | Abietadienol; Abietadienal; Levopimaradienol; Isopimara-7,15-dienol; Isopimara-7,15-dienal; Dehydroabietadienol; Dehydroabietadienal | N/A | N/A | 40 |
| Q9FRK4 | CYP721A1 | CYP | *Arabidopsis thaliana* | N/A | N/A | N/A | 40 |
| F4HP86 | CYP722A1 | CYP | *Arabidopsis thaliana* | N/A | N/A | N/A | 40 |
| Q6F4F5 | CYP724B1 | CYP | *Oryza sativa subsp. japonica* | Brassinosteroid; 22-OHCR; 22-hydroxyCR | Anti-cancer; Anti-inflammatory | 15705958; 16369540; 35253313; 31400389 | 40 |
| Q9AXM6 | CYP725A1 | CYP | *Taxus cuspidata* | Taxadien-5-alpha-acetoxy-10-beta-ol | Chemotherapeutic lead compound | 11396929; 21160477 | 40 |
| B9RHW2 | CYP726A13 | CYP | *Ricinus communis* | Ricinoleic acid | Bactericidal; Anti-inflammatory; Antiherpetic agents | 30733060; 35661063; 25542985 | 40 |
| A0A0G7ZP21 | CYP727C1 | CYP | *Picea glauca* | N/A | N/A | N/A | 40 |
| A0A291FAT9 | CYP728S3 | CYP | *Taxus wallichiana var. chinensis* | N/A | N/A | N/A | 40 |
| A0A291FAZ9 | CYP729B25 | CYP | *Taxus wallichiana var. chinensis* | N/A | N/A | N/A | 40 |
| A0A7L7RB92 | CYP733A1 | CYP | *Nothapodytes nimmoniana* | N/A | N/A | N/A | 40 |
| O48786 | CYP734A1 | CYP | *Arabidopsis thaliana* | N/A | N/A | N/A | 40 |
| Q9ZW95 | CYP735A2 | CYP | *Arabidopsis thaliana* | Trans-zeatin | Antioxidant | 36009800 | 40 |
| A0A068Q6L2 | CYP736A117 | CYP | *Prunus mume* | N/A | N/A | N/A | 40 |
| O82381 | UGT71C1 | UGT | *Arabidopsis thaliana* | Trans-resveratrol; Curcumin, Vanillin and Etoposide | Anti-aging; Antioxidant; Anti-inflammatory; Anti-cancer; Cancer treatment; Neuroprotective agent | 27058985; 29210129; 33182828; 34754179; 34610223; 34202987 | 45 |
| Q9LVR1 | UGT72E2 | UGT | *Arabidopsis thaliana* | Monolignols; Coniferyl alcohol 4-O-glucoside | Antioxidant | 24667164; 21149736; 16995900; 15907484; 11042211; 12903947 | 45 |
| Q7Y232 | UGT73B4 | UGT | *Arabidopsis thaliana* | 2-HADNT-O-monoglucoside | Bioremediation | 18702669 | 45 |
| O22822 | UGT74F2 | UGT | *Arabidopsis thaliana* | Salicylic Acid glucoside | Anti-inflammatory | 27718358; 28425481 | 45 |
| Q9LR44 | UGT75B1 | UGT | *Arabidopsis thaliana* | 4-aminobenzoate glucose ester | Abiotic stress response in Plants | 31894456 | 45 |
| Q6VAB4 | UGT76G1 | UGT | *Stevia rebaudiana* | Steviol glycosides | Sweetener; Anti-hypertensive; Anti-obesity; Anti-diabetic; Antioxidant; Anti-cancer; Antimicrobial effects | 30569490; 15610349; 34803554 | 45 |
| A0A2Z5CVA1 | UGT77B2 | UGT | *Crocosmia x crocosmiiflora* | Montbretin A | Drug candidate to treat type-2 diabetes | 29967287; 31004005; 29967287 | 45 |
| A6XNC6 | UGT78G1 | UGT | *Medicago truncatula* | Anthocyanin; Kaempferol glycosides; Isoquercetin | Antioxidant; Antioxidant; Anti-inflammatory | 19683002; 36557809; 30817903; 19898800 | 45 |
| Q9LVW3 | UGT79B1 | UGT | *Arabidopsis thaliana* | Anthocyanin | Antioxidant | 21899608; 34388293; 36557809 | 45 |
| Q9M8Z7 | UGT80A2 | UGT | *Arabidopsis thaliana* | Steryl glycosides; Acyl steryl glycosides | Anti-inflammatory; Anti-diabetic; Anti-microbial | 32084504; 34685527; 12403168 | 45 |
| O81770 | UGT81A1 | UGT | *Arabidopsis thaliana* | Galactolipid | Anti-inflammatory | 11171188; 10869420; 11171188; 16253232 | 45 |
| Q9LHJ2 | UGT82A1 | UGT | *Arabidopsis thaliana* | N/A | N/A | N/A | 45 |
| Q9SGA8 | UGT83A1 | UGT | *Arabidopsis thaliana* | Sinapoyl anthocyanin | Antioxidant | 32420359; 25976793 | 45 |
| Q5XF20 | UGT84A1 | UGT | *Arabidopsis thaliana* | Glucosylate the phytotoxic xenobiotic compound 2,4,5-trichlorophenol | Bioremediation | 11042211; 11187886; 12721858 | 45 |
| Q9SBL1 | UGT85B1 | UGT | *Sorghum bicolor* | Dhurrin | Natural pest control agent | 17706731; 26493517; 36279963 | 45 |
| Q9ZUV0 | UGT86A2 | UGT | *Arabidopsis thaliana* | N/A | N/A | N/A | 45 |
| O64733 | UGT87A2 | UGT | *Arabidopsis thaliana* | N/A | N/A | 27747895 | 45 |
| A6BM07 | UGT88E3 | UGT | *Glycine max* | Isoflavonoids | Anti-atherosclerotic; Alleviating Menopausal Symptoms; Anti-cancer | 26019269; 31936113; 12798527; 32059369 | 45 |
| Q9LNE6 | UGT89C1 | UGT | *Arabidopsis thaliana* | Quercetin 7-O-rhamnoside; Kaempferol 7-O-rhamnoside | Anti-inflammatory; Antioxidant | 22264152; 36076438 | 45 |
| Q9ZVX4 | UGT90A1 | UGT | *Arabidopsis thaliana* | N/A | N/A | 35628539 | 45 |
| B3VI56 | UGT91D2 | UGT | *Stevia rebaudiana* | Steviol glycosides | Sweetener; Anti-hypertensive; Anti-obesity; Anti-diabetic; Antioxidant; Anti-cancer; Antimicrobial effects | 27923373; 26358188; 34803554 | 45 |
| Q9LXV0 | UGT92A1 | UGT | *Arabidopsis thaliana* | N/A | N/A | N/A | 45 |
| A0A3Q9EMT7 | UGT93B8 | UGT | *Avena strigosa* | N/A | N/A | N/A | 45 |
| Q5NTH0 | UGT94B1 | UGT | *Bellis perennis* | Glucuronosylated anthocyanins; Ginsenoside Rd | Antioxidant; Neuroprotective agent | 28315473; 18829982; 34952508; 12358498; 15509561 | 45 |
| A0A067XTB0 | UGT95B1 | UGT | *Cicer arietinum* | N/A | N/A | N/A | 45 |
| I2BHE9 | UGT97B3 | UGT | *Linum usitatissimum* | N/A | N/A | N/A | 45 |
| A0A3S5HT30 | UGT98B4 | UGT | *Avena strigosa* | N/A | N/A | N/A | 45 |
| A0A3S9LYS9 | UGT99A6 | UGT | *Avena strigosa* | N/A | N/A | N/A | 45 |
| A0A0K0PVW1 | UGT100 | UGT | *Panax ginseng* | Protopanaxatriol | Anti-cancer; Neurogenesis in Alzheimer's disease | 26032089; 27746309; 27446225; 37067038 | 45 |
| Q9SXD9 | GSXL7 | FMO | *Arabidopsis thaliana* | Glucoraphanin | Anti-inflammatory | 29898626; 29898626; 31713870 | 35 |
| Q9FKE7 | FMO2 | FMO | *Arabidopsis thaliana* | Indole-3-pyruvic acid | Anti-inflammatory | 30429284; 30420566 | 35 |
| O49312 | YUC7 | FMO | *Arabidopsis thaliana* | Indole-3-acetic acid | Anti-inflammatory | 22025724; 31484323; 35185872 | 35 |
| O49499 | CAMT4 | OMT | *Arabidopsis thaliana* | Ferulic Acid; Scopoletin | Alzheimer's Therapeutic; Anti-cancer | 34963433; 27457122; 33931140; 33292145 | 30 |
| Q6XMI3 | BSMT1 | OMT | *Arabidopsis thaliana* | Methylsalicylate | Analgesic | 18987162; 20488836 | 30 |
| Q9FYC4 | FAMT | OMT | *Arabidopsis thaliana* | Methyl farnesoate | Natural pest control agent | 16165084; 32849271 | 30 |
| Q9C9W4 | TSM1 | OMT | *Arabidopsis thaliana* | Phenylpropanoid polyamine conjugate | Anti-microbial | 20800856; 18557837 | 30 |
| Q9LPU6 | IGMT3 | OMT | *Arabidopsis thaliana* | Indole glucosinolate | Anti-inflammatory | 24360830; 27810943 | 30 |
| Q9SCP7 | NAMT1 | OMT | *Arabidopsis thaliana* | Niacin | Neuroprotective agent; Prevention of cardiovascular disease | 34171402; 28533213; 22646128; 28616955 | 30 |
| A8QW52 | OMT1 | OMT | *Sorghum bicolor* | Methyleugenol | Anti-inflammatory | 24420851; 30634080; 20815773; 34358859 | 30 |
| Q9C8E3 | TPS26 | TPS | *Arabidopsis thaliana* | N/A | N/A | 27933080 | 30 |
| A0A178U9Y5 | TP20L | TPS | *Arabidopsis thaliana* | Dolabella-3,7-dien-18-ol | Anti-tumor | 27933080; 36499119 | 30 |
| Q84UV0 | AtTPS14 | TPS | *Arabidopsis thaliana* | (S)-linalool | Anti-Parkinsonian's effect | 35665972; 12566586; 35665972 | 30 |
| Q9SHG0 | TPS3 | TPS | *Arabidopsis thaliana* | N/A | N/A | N/A | 30 |
| Q9T079 | TPS4 | TPS | *Arabidopsis thaliana* | N/A | N/A | N/A | 30 |
| G5CV46 | TPS31 | TPS | *Solanum lycopersicum* | Viridiflorene | N/A | 21818683 | 30 |
| G5CV37 | G5CV37 | TPS | *Solanum lycopersicum* | N/A | N/A | N/A | 30 |
| G9MAN7 | smTPS4 | TPS | *Selaginella moellendorffii* | (+)-copalyl diphosphate and miltiradiene | N/A | 22027823 | 30 |
| Q39103 | DOXC3 | DOXC | *Arabidopsis thaliana* | GA1; GA4 | N/A | 9625708 | 40 |
| Q4PT02 | DOXC7 | DOXC | *Arabidopsis thaliana* | GA53; GA20 | N/A | N/A | 40 |
| Q9FZ21 | DOXC12 | DOXC | *Arabidopsis thaliana* | GA51; GA29; GA34; GA8 | N/A | N/A | 40 |
| Q9C6I4 | DOXC13 | DOXC | *Arabidopsis thaliana* | N/A | N/A | N/A | 40 |
| F4JAD4 | DOXC14 | DOXC | *Arabidopsis thaliana* | N/A | N/A | N/A | 40 |
| Q9XI76 | DOXC15 | DOXC | *Arabidopsis thaliana* | N/A | N/A | N/A | 40 |
| O65485 | DOXC17 | DOXC | *Arabidopsis thaliana* | N/A | N/A | N/A | 40 |
| Q8VYD7 | DOXC19 | DOXC | *Arabidopsis thaliana* | N/A | N/A | N/A | 40 |
| Q9M9D9 | DOXC20 | DOXC | *Arabidopsis thaliana* | N/A | N/A | N/A | 40 |
| Q9LJ66 | DOXC21 | DOXC | *Arabidopsis thaliana* | N/A | N/A | N/A | 40 |
| Q8H113 | DOXC23 | DOXC | *Arabidopsis thaliana* | N/A | N/A | N/A | 40 |
| A1A6I8 | DOXC24 | DOXC | *Arabidopsis thaliana* | N/A | N/A | N/A | 40 |
| Q9C6F0 | DOXC27 | DOXC | *Arabidopsis thaliana* | N/A | N/A | N/A | 40 |
| Q9S818 | DOXC28 | DOXC | *Arabidopsis thaliana* | Anthocyanins | Anti-cancer; Anti-inflammatory; Antioxidant | 28974032; 28799785; 36388502; 31141884 | 40 |
| Q9C899 | DOXC30 | DOXC | *Arabidopsis thaliana* | Scopoletin | Anti-cancer; Antioxidant | 33292145; 18547395; 35498835 | 40 |
| Q94A78 | DOXC31 | DOXC | *Arabidopsis thaliana* | N/A | N/A | N/A | 40 |
| Q9LTH7 | DOXC31 | DOXC | *Arabidopsis thaliana* | N/A | N/A | N/A | 40 |
| F4INZ9 | DOXC37 | DOXC | *Arabidopsis thaliana* | N/A | N/A | N/A | 40 |
| Q9ZSA8 | DOXC38 | DOXC | *Arabidopsis thaliana* | 2,3-dihydroxybenzoic acid | Anti-microbial | 24449781; 23959884 | 40 |
| F4JLS2 | DOXC45 | DOXC | *Arabidopsis thaliana* | N/A | N/A | N/A | 40 |
| O80449 | DOXC46 | DOXC | *Arabidopsis thaliana* | 12-hydroxyjasmonate | N/A | 28559313 | 40 |
| Q96323 | DOXC47 | DOXC | *Arabidopsis thaliana* | Anthocyanin | Anti-cancer; Anti-inflammatory; Antioxidant | 28974032; 28799785; 31141884; 12940955 | 40 |
| Q9FFQ4 | DOXC47 | DOXC | *Arabidopsis thaliana* | N/A | N/A | N/A | 40 |
| Q9SYM7 | DOXC52 | DOXC | *Arabidopsis thaliana* | N/A | N/A | N/A | 40 |
| Q06588 | DOXC53 | DOXC | *Arabidopsis thaliana* | Ethylene | N/A | 17993622 | 40 |
| Q9LIF4 | DOXC54 | DOXC | *Arabidopsis thaliana* | N/A | N/A | N/A | 40 |
| Q9FN27 | DOXC55 | DOXC | *Arabidopsis thaliana* | N/A | N/A | N/A | 40 |
| P43309 | MdPPO | PPO | *Malus domestica* | 1,2-benzoquinone | N/A | 19783226 | 40 |
| AFW89544 | COX10 | PT | *Zea mays* | N/A | N/A | N/A | 35 |
| ABB70123 | HPT_VTE2-1 | PT | *Triticum aestivum* | Tocopherol | Anti-neurodegenerative Drugs; Potential Therapeutics for Stress-Induced Disorders; Antioxidant; Anti-cancer | 15377170; 21223386; 29173163; 32017273 | 35 |
| BAP27988 | ClPT1 | PT | *Citrus limon* | 8-geranylumbelliferone | Anti-Inflammatory; Cancer Therapeutics | 25077796; 34411570; 25077796 | 35 |
| BAG12671 | N8DT-1 | PT | *Sophora flavescens* | Sophoraflavanone G | Anti-cancer; Anti-inflammatory | 35682783; 35426453; 18218974 | 35 |
| Q0WUA3 | ABC4 | PT | *Arabidopsis thaliana* | Phylloquinone | Nutritional supplements; Therapeutics for Cardiovascular Diseases | 34472618; 15686525 | 35 |
| Q38833 | CHLG | PT | *Arabidopsis thaliana* | Chlorophyll a | Antioxidant; Anti-cancer | 32828967; 25041167 | 35 |
| Q93YP7 | PPT1 | PT | *Arabidopsis thaliana* | Coenzyme Q | Energy supplements; Antioxidant; Neuroprotective Agents; Treatment for cardiovascular disease | 22161420; 24389208; 15604701; 34067632 | 35 |
| F4J8K0 | HST | PT | *Arabidopsis thaliana* | Plastoquinone-9; Tocochromanol | Anti-aging; Antioxidant; Neuroprotective agent; Anti-cancer | 19159610; 16989822; 20385544 | 35 |
| BAO31627 | PcPT | PT | *Petroselinum crispum* | Demethylsuberosin | Anti-inflammatory | 31635294**;** 24354545 | 35 |
| BBG56301 | AcPT1 | PT | *Artemisia capillaris* | Artepillin C | Antioxidant; Anti-inflammatory; Anti-microbial; Anti-diabetic; Anti-tumor; Neuroprotective; Gastroprotective; Immunomodulatory effects | 33152464; 31646187 | 35 |
| BBD96134 | RdPT1 | PT | *Rhododendron dauricum* | Daurichromenic Acid | Anti-HIV | 14602030; 30097469 | 35 |
| AAP43911 | HGGT | PT | *Hordeum vulgare* | Tocotrienol | Antioxidant; Anti-inflammatory; Anti-cancer | 32017273; 12897790 | 35 |
| BAJ61049 | PT1 | PT | *Humulus lupulus* | Beta-bitter acid | Anti-inflammatory; Anti-cancer; Anti-microbial; 32269899 | 18162387; 24954859; 22166201 | 35 |
| AZK16227 | Ptpat | PT | *Hypericum calycinum* | Patulone | Treatment of PAF-mediated diseases, such as allergy, inflammation, thrombosis and asthma | 30485455; 15678383 | 35 |
| AJD80983 | FPT | PT | *Maclura tricuspidata* | Cudraflavanone | Neuroprotective agent; Anti-inflammatory | 25361766; 26907256 | 35 |
| BAB84122 | PGT-1 | PT | *Lithospermum erythrorhizon* | Shikonin | Anti-cancer; Protective effect against skin diseases; Anti-inflammatory | 19392660; 11744717; 31553936 | 35 |
| BAF73620 | GmIF7MaT (GmAT133) | BAHD | *Glycine max* | Isoflavone 7-O-(6″-O-malonyl-β-d-glucosides) | N/A | 17602715 | 30 |
| AAL50565 | Pf5MaT | BAHD | *Perilla frutescens* | Malonylated anthocyanin | Anti-cancer; Anti-inflammatory; Antioxidant | 28974032; 28799785; 31141884 | 30 |
| NP_171890 | At3AT1 | BAHD | *Arabidopsis thaliana* | Coumaroylated anthocyanin | Anti-cancer; Anti-inflammatory; Antioxidant | 28974032; 28799785; 31141884 | 30 |
| CAA61258 | ZmGlossy2 | BAHD | *Zea mays* | N/A | N/A | 8580961 | 30 |
| AAM64817 | CER2 | BAHD | *Arabidopsis thaliana* | Epicuticular wax | N/A | 17376164; 22930748 | 30 |
| AAC99311 | CrDAT | BAHD | *Catharanthus roseus* | Vindoline | Anti-tumor; Anti-diabetic | 2350183; 11154328; 24012527; 25918424 | 30 |
| AAG13130 | FaSAAT | BAHD | *Fragaria x ananassa* | Esters | Perfumes and Flavors; Pharmaceuticals; Cosmetics; Green solvents; Advanced biofuels | 31986468; 10810141 | 30 |
| AAK73661 | PsSalAT | BAHD | *Papaver somniferum* | Thebaine; Morphine | Analgesic; Pharmaceutical synthesis ingredient | 22098111; 22098111; 9061094 | 30 |
| ABG75942 | PhCFAT | BAHD | *Petunia x hybrida* | Coniferyl acetate; Isoeugenol | Antibacterial | 17241449; 17241449; 17241449; 35367366 | 30 |
| AAO73071 | HvACT | BAHD | *Hordeum vulgare* | Hordatines | Anti-fungal | 12582168; metabolites | 30 |
| Q94CD1 | HHT1 | BAHD | *Arabidopsis thaliana* | Suberin | Anti-microbial | 19846769; 19759341; 35155893; 35030736 | 30 |
| AAL92459 | TcBAPT | BAHD | *Taxus cuspidata* | Taxol | Cancer treatment | 12232048; 12232048; 25213191 | 30 |
| BAD89275 | LaHMT/HLT | BAHD | *Lupinus albus* | Lupanine alkaloids | Potential Therapeutic Agent for Diabetes; Anthelmintic | 15659437; 8195240; 26492234; 31227784 | 30 |
| ACI16631 | TpHCT2 | BAHD | *Trifolium pratense* | Phaselic acid | N/A | 19525325 | 30 |
| NP_199704 | AtHCT | BAHD | *Arabidopsis thaliana* | Lignin | Biofuel | 15161961; 26858288; 27805809; 25051990 | 30 |

| Uniprot/NCBI ID | Enzyme | Family | Organism | Associated Compound | Potential use | PMID | Blast Identity threshold |
| --- | --- | --- | --- | --- | --- | --- | --- |
| A0A3Q9R4N5 | CYP76B74 | CYP | *Arnebia euchroma* | Shikonin | Anti-cancer; Protective effect against skin diseases; Anti-inflammatory | 30498024; 30498024; 11744717; 31553936 | 55 |
| QUD18410 | CYP82AR | CYP | *Lithospermum erythrorhizon* | Shikonin | Anti-cancer; Protective effect against skin diseases; Anti-inflammatory | 33728922; 11744717; 31553936 | 55 |
| BBV14785 | LeSAT | BAHD | *Lithospermum erythrorhizon* | Shikonin | Anti-cancer; Protective effect against skin diseases; Anti-inflammatory | 32727911; 11744717; 31553936 | 60 |
| BAB84122 | PGT-1 | PT | *Lithospermum erythrorhizon* | Shikonin | Anti-cancer; Protective effect against skin diseases; Anti-inflammatory | 19392660; 11744717; 31553936 | 60 |
| BAB08004 | 6-OMT | OMT | *Coptis japonica* | Benzylisoquinoline | Antidepressant; Neurological disorders treatment; Analgesic | 19601791; 10811648; 33340158 | 50 |
| BAB71802 | CNMT | NMT | *Coptis japonica* | Benzylisoquinoline | Antidepressant; Neurological disorders treatment; Analgesic | 19601791; 11682473; 33340158 | 50 |
| BAB08005 | 4-OMT | OMT | *Coptis japonica* | Benzylisoquinoline | Antidepressant; Neurological disorders treatment; Analgesic | 19601791; 10811648; 33340158 | 50 |

**Table S2**. Comprehensive enzyme information associated to shikonin and benzylisoquinoline biosynthesis with emphasis on functions, sequences, and blast identity thresholds

**Table S3.** Environmental data used for distribution prediction by MaxEnt

| No | Category | Variable |
| --- | --- | --- |
| 1 | Bioclimatic data | Annual Mean Temperature |
| 2 |  | Mean diurnal range (mean of monthly (max temp - min temp)) |
| 3 |  | Isothermality |
| 4 |  | Temperature seasonality |
| 5 |  | Max temperature of warmest month |
| 6 |  | Min temperature of coldest month |
| 7 |  | Temperature annual range |
| 8 |  | Mean temperature of wettest quarter |
| 9 |  | Mean temperature of driest quarter |
| 10 |  | Mean temperature of warmest quarter |
| 11 |  | Mean temperature of coldest quarter |
| 12 |  | Annual precipitation |
| 13 |  | Precipitation of wettest month |
| 14 |  | Precipitation of driest month |
| 15 |  | Precipitation seasonality |
| 16 |  | Precipitation of wettest quarter |
| 17 |  | Precipitation of driest quarter |
| 18 |  | Precipitation of warmest quarter |
| 19 |  | Precipitation of coldest quarter |
| 20 | Solar Radiation | Annual total solar radiation (kJ m-2 day-1) |
| 21 | Water vapor pressure | Annual average water vapor pressure (kPa) |
| 22 |  | Minimum monthly average water vapor pressure (kPa) |
| 23 |  | Maximum monthly average water vapor pressure (kPa) |
| 24 | Wind speed | Annual average wind speed (m s-1) |
| 25 |  | Minimum monthly average wind speed (m s-1) |
| 26 |  | Maximum monthly average wind speed (m s-1) |
| 27 | Last glacial maximum bioclimatic data | Annual mean temperature |
| 28 |  | Mean diurnal range (mean of monthly (max temp - min temp)) |
| 29 |  | Isothermality |
| 30 |  | Temperature seasonality |
| 31 |  | Max temperature of warmest month |
| 32 |  | Min temperature of coldest month |
| 33 |  | Temperature annual range |
| 34 |  | Mean temperature of wettest quarter |
| 35 |  | Mean temperature of driest quarter |
| 36 |  | Mean temperature of warmest quarter |
| 37 |  | Mean temperature of coldest quarter |
| 38 |  | Annual precipitation |
| 39 |  | Precipitation of wettest month |
| 40 |  | Precipitation of driest month |
| 41 |  | Precipitation seasonality |
| 42 |  | Precipitation of wettest quarter |
| 43 |  | Precipitation of driest quarter |
| 44 |  | Precipitation of warmest quarter |
| 45 |  | Precipitation of coldest quarter |
| 46 | Terrain | Maximum elevation (m) |
| 47 |  | Average elevation (m) |
| 48 |  | Minimum elevation (m) |
| 49 |  | Standard deviation of elevation |
| 50 | Land area | Land area (km^2) |
| 51 |  | Land area ratio (%) |
| 52 | Land use area | City area (km^2) |
| 53 |  | Farmland area (km^2) |
| 54 |  | Forest area (km^2) |
| 55 |  | Grassland area (km^2) |
| 56 | Soil factor | Depth to bedrock (r horizon) if observed, depth to bedrock (m) |
| 57 |  | Bulk density in tons per cubic-meter, bulk density (tons/cubic meter) |
| 58 |  | Volume percentage of coarse fragments (> 2 mm), volume percentage of surface gravel (> 2 mm) |
| 59 |  | Sand content in percent, sand content percentage in the surface layer |
| 60 |  | Salt content in percent, surface layer silt content percentage |
| 61 |  | Clay content in percent, percentage of clay content in the surface layer |
| 62 |  | Cation exchange capacity in cmol per kilogram, surface layer cation exchange capacity (cmol/kg) |
| 63 |  | Soil ph in water suspension, ph of surface soil suspension |
| 64 |  | Moisture potential in kpa e.g., -10 (pF 2.0), surface moisture potential (kPa) |
| 65 |  | Soil organic carbon concentration in permille or g / kg, surface soil organic carbon concentration (permille) |

**Figure S1.** The completeness of the genetic and distributional data. (A) Predicted SR from species possessing genomic or transcriptomic data (B) The proportion of species possessing genetic data in relation to predicted SR. (C) Plots showing the relationship between predicted SR and predicted SR from species possessing genomic or transcriptomic data of each cell.
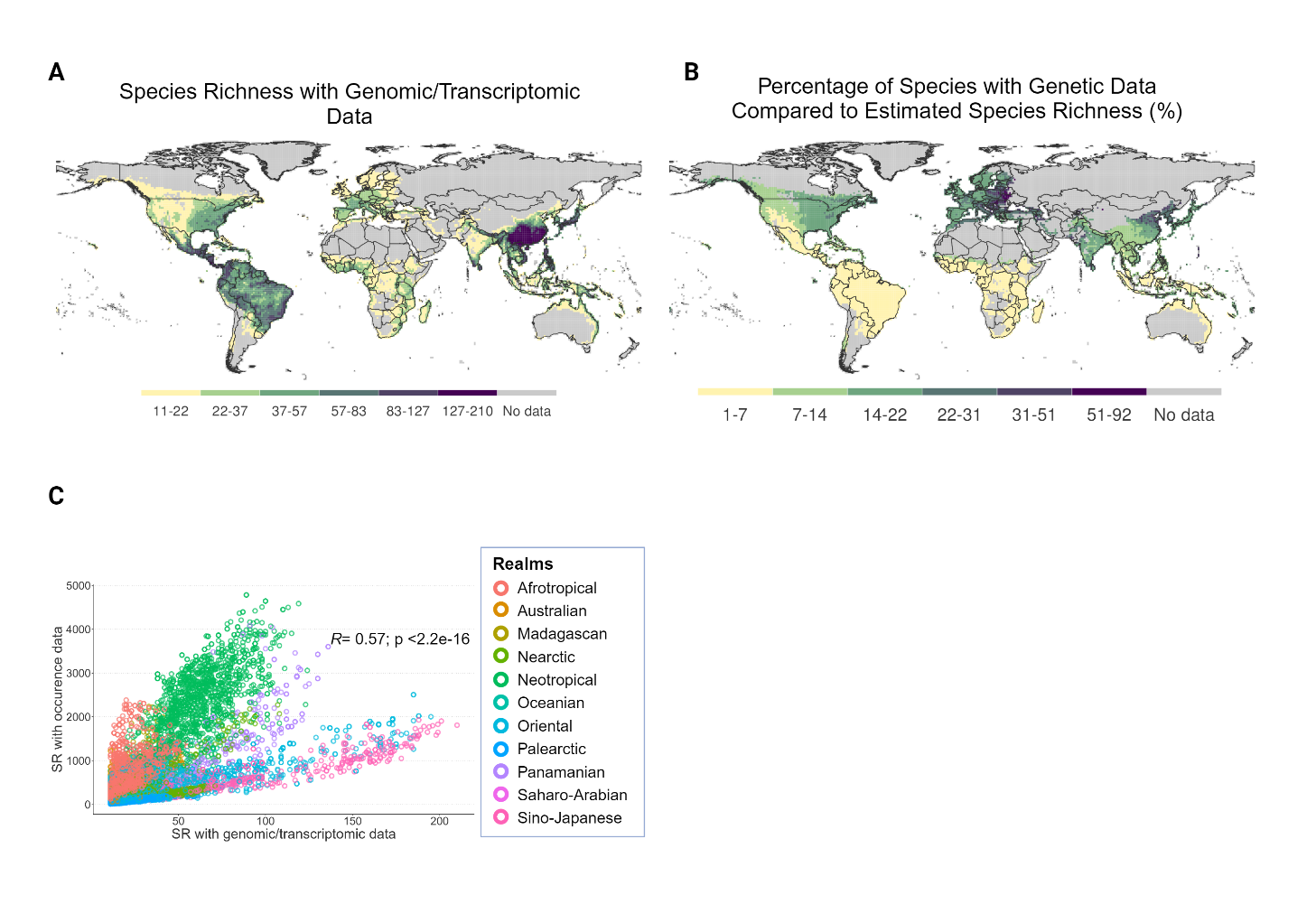

**Figure S2.** Global pattern of (A) GmIF7MaT, (B) Pf5MaT, (C) At3AT1, (D) ZmGlossy2, (E) CER2, (F), CrDAT, (G) FaSAAT, (H) PsSalAT, (I) PhCFAT, and (J) HvACT

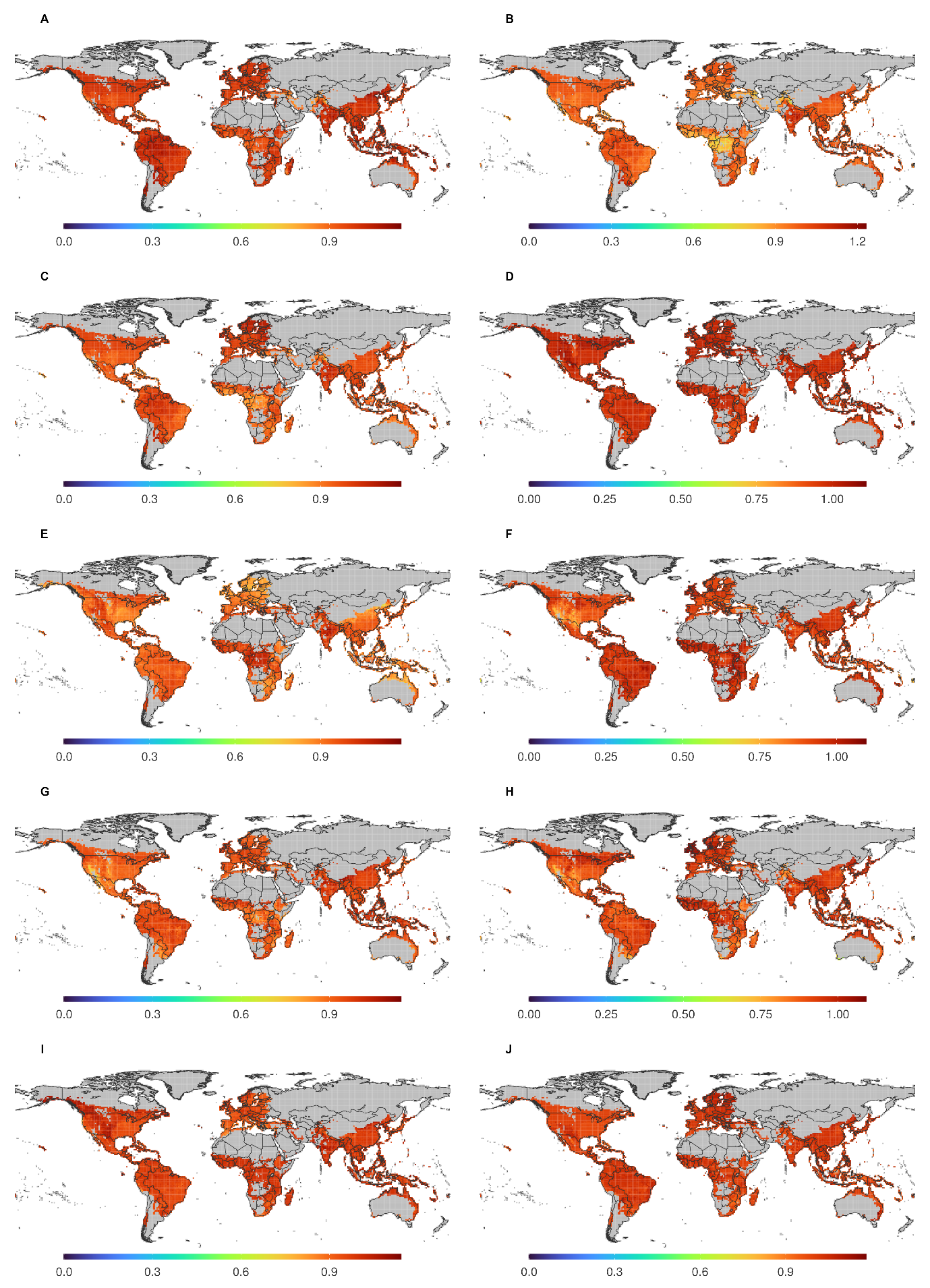

**Figure S3.** Global pattern of (A) HHT1, (B) TcBAPT, (C) LaHMT/HLT, (D) TpHCT2, (E) AtHCT, (F) DOXC3, (G) DOXC7, (H) DOXC12, (I) DOXC13, and (J) DOXC14

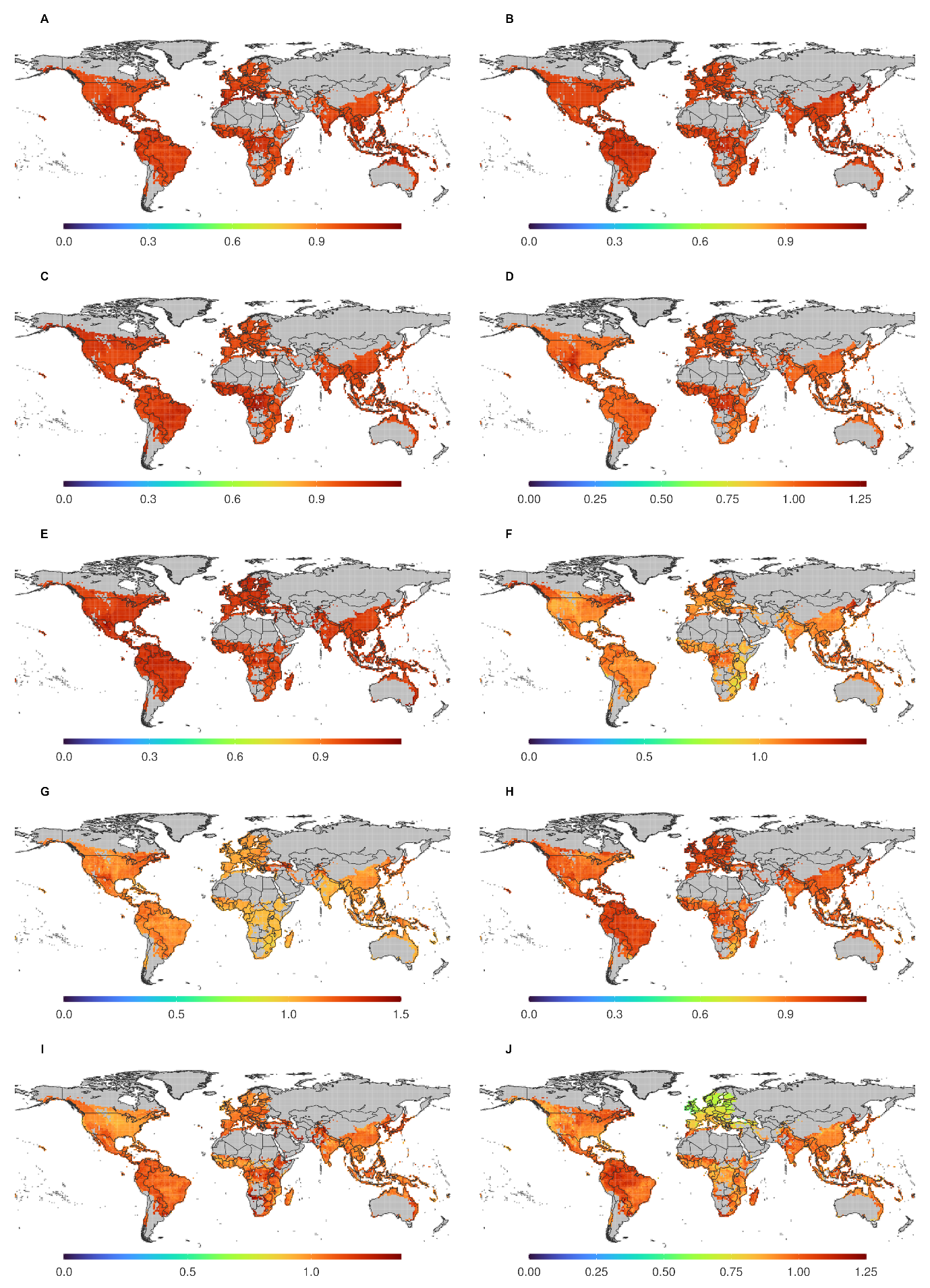

**Figure S4.** Global pattern of (A) DOXC15, (B) DOXC17, (C) DOXC19, (D) DOXC20, (E) DOXC21, (F) DOXC23, (G) DOXC24, (H) DOXC27, (I) DOXC28, and (J) DOXC30

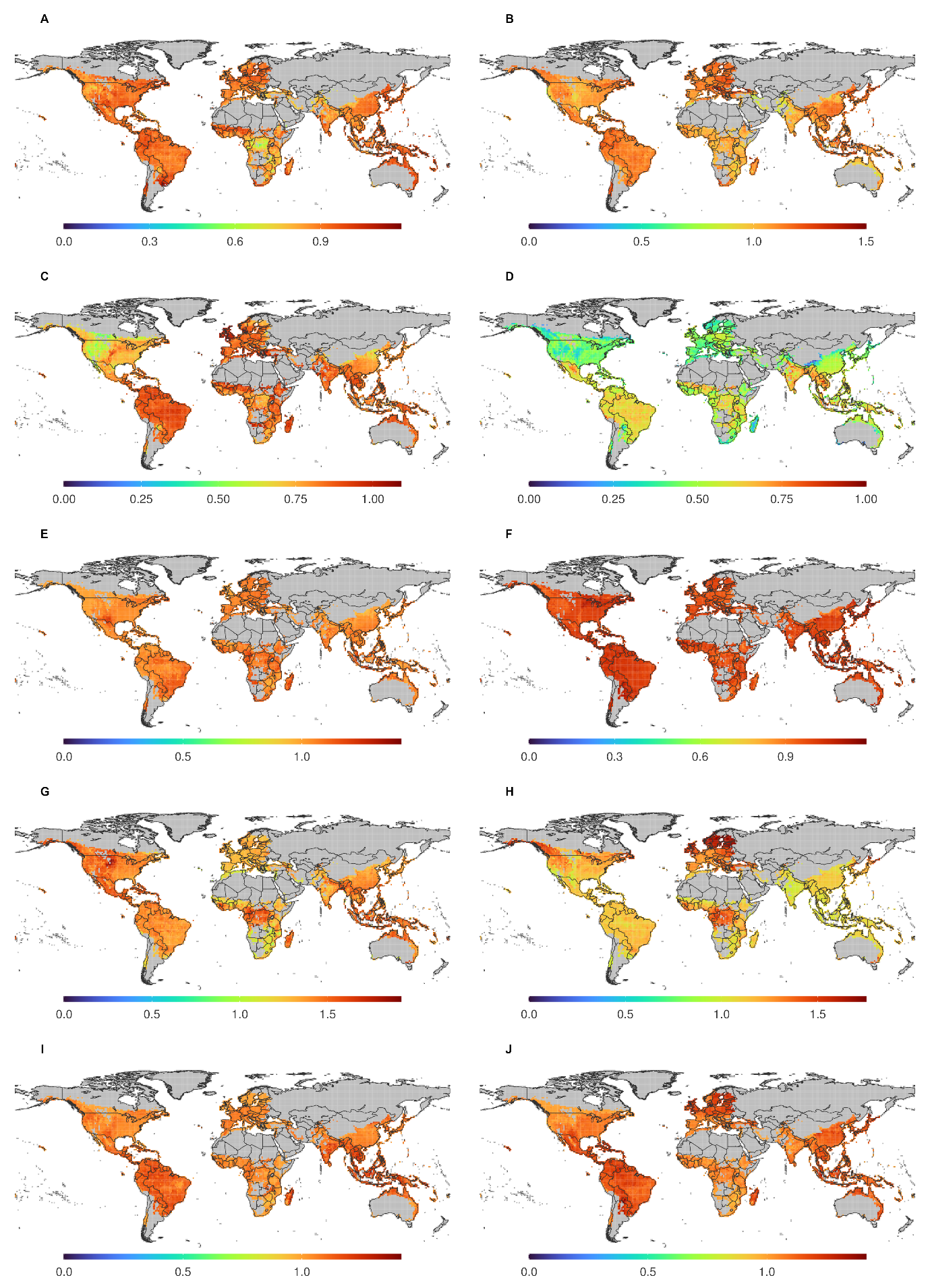

**Figure S5.** Global pattern of (A) DOXC31, (B) DOXC31, (C) DOXC37, (D) DOXC38, (E) DOXC45, (F) DOXC46, (G) DOXC47, (H) DOXC47, (I) DOXC52, and (J) DOXC53

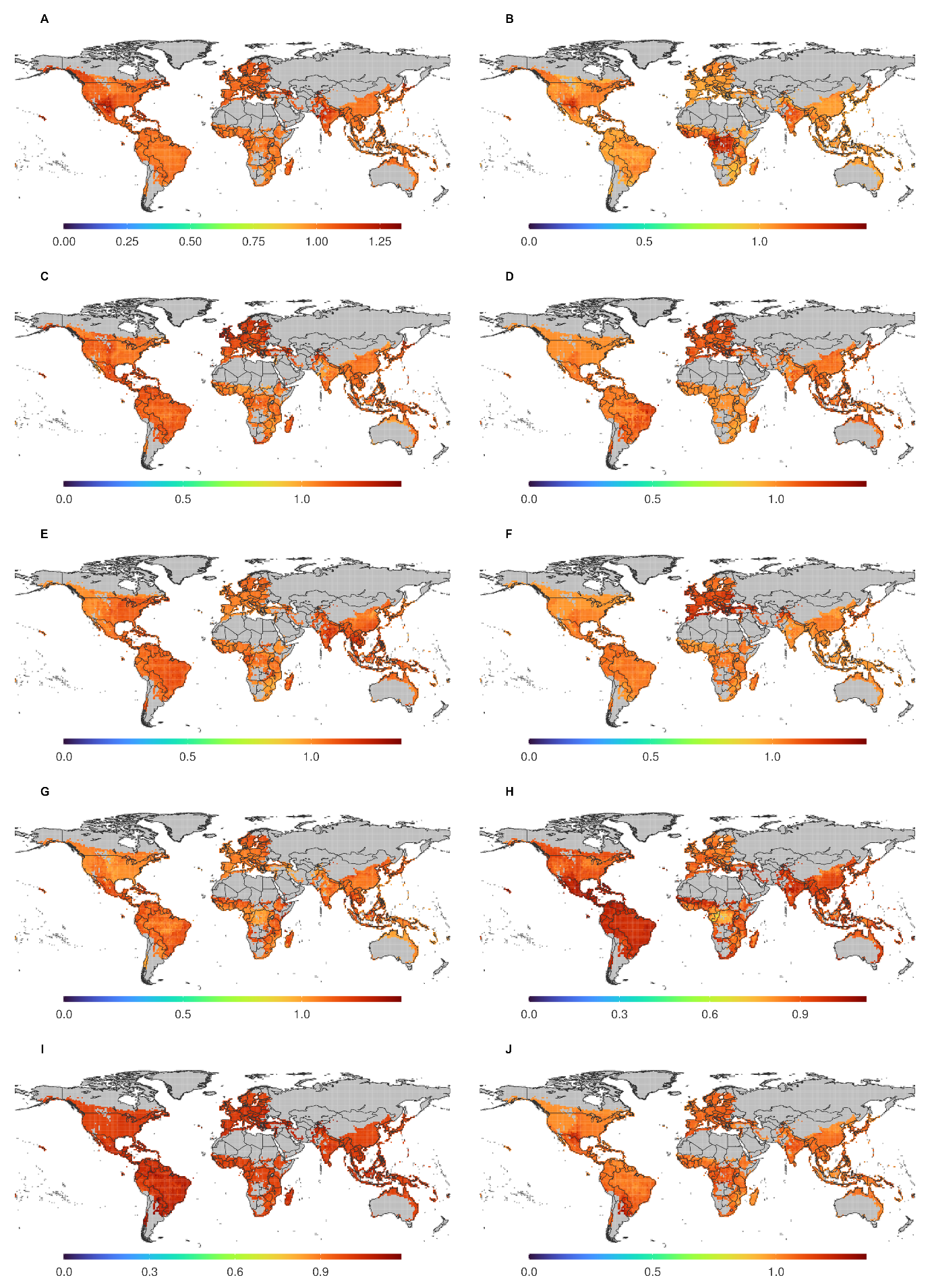

**Figure S6.** Global pattern of (A) DOXC54, (B) DOXC55, (C) GSXL7, (D) FMO2, (E) YUC7, (F) CAMT4, (G) BSMT1, (H) FAMT, (I) TSM1, and (J) IGMT3

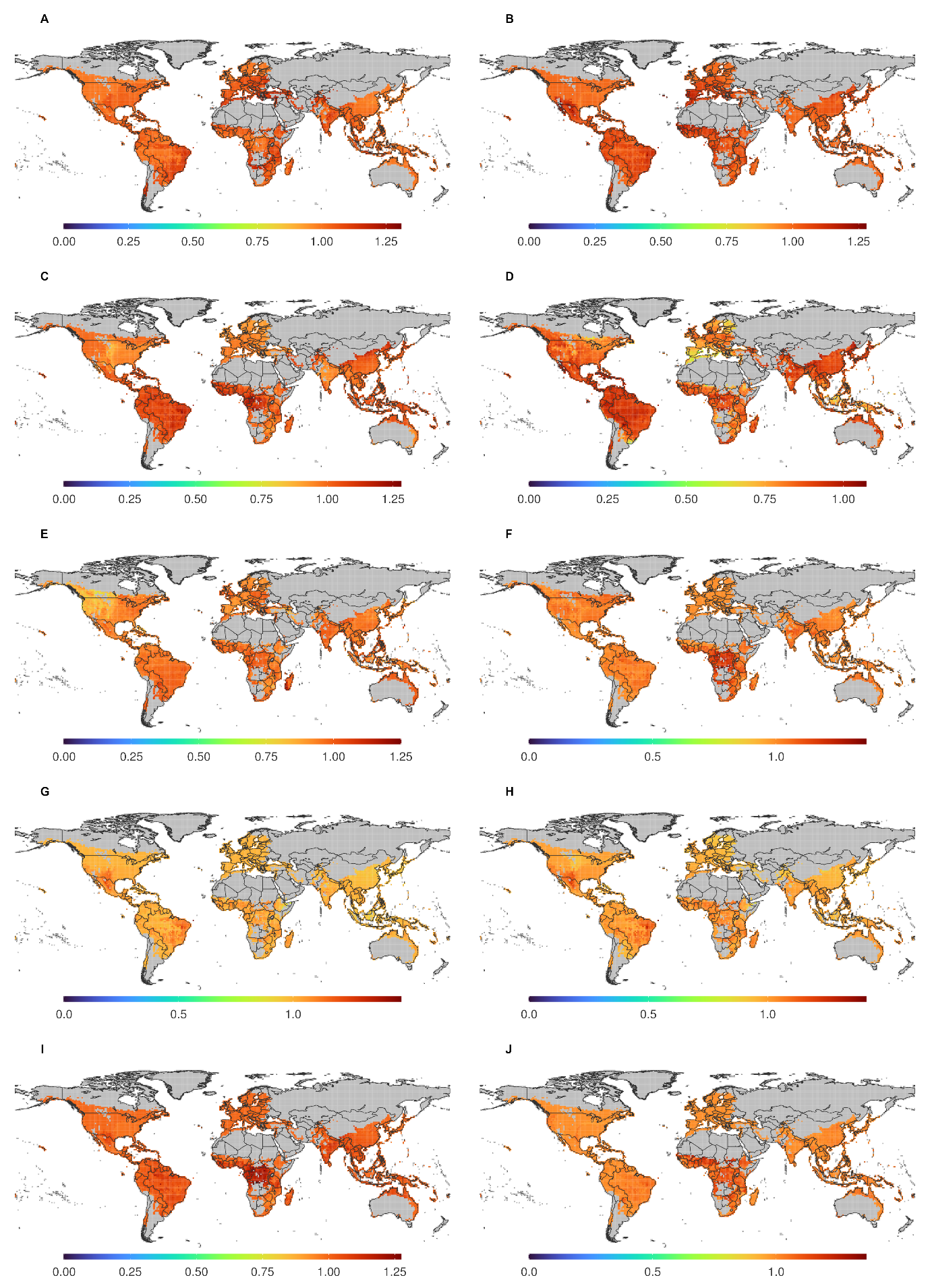

**Figure S7.** Global pattern of (A) NAMT1, (B) OMT1, (C) CYP51G1, (D) CYP71D1V2, (E) CYP72A1, (F) CYP73A1, (G) CYP74D2, (H) CYP75B2, (I) CYP76M8, and (J) CYP77A2

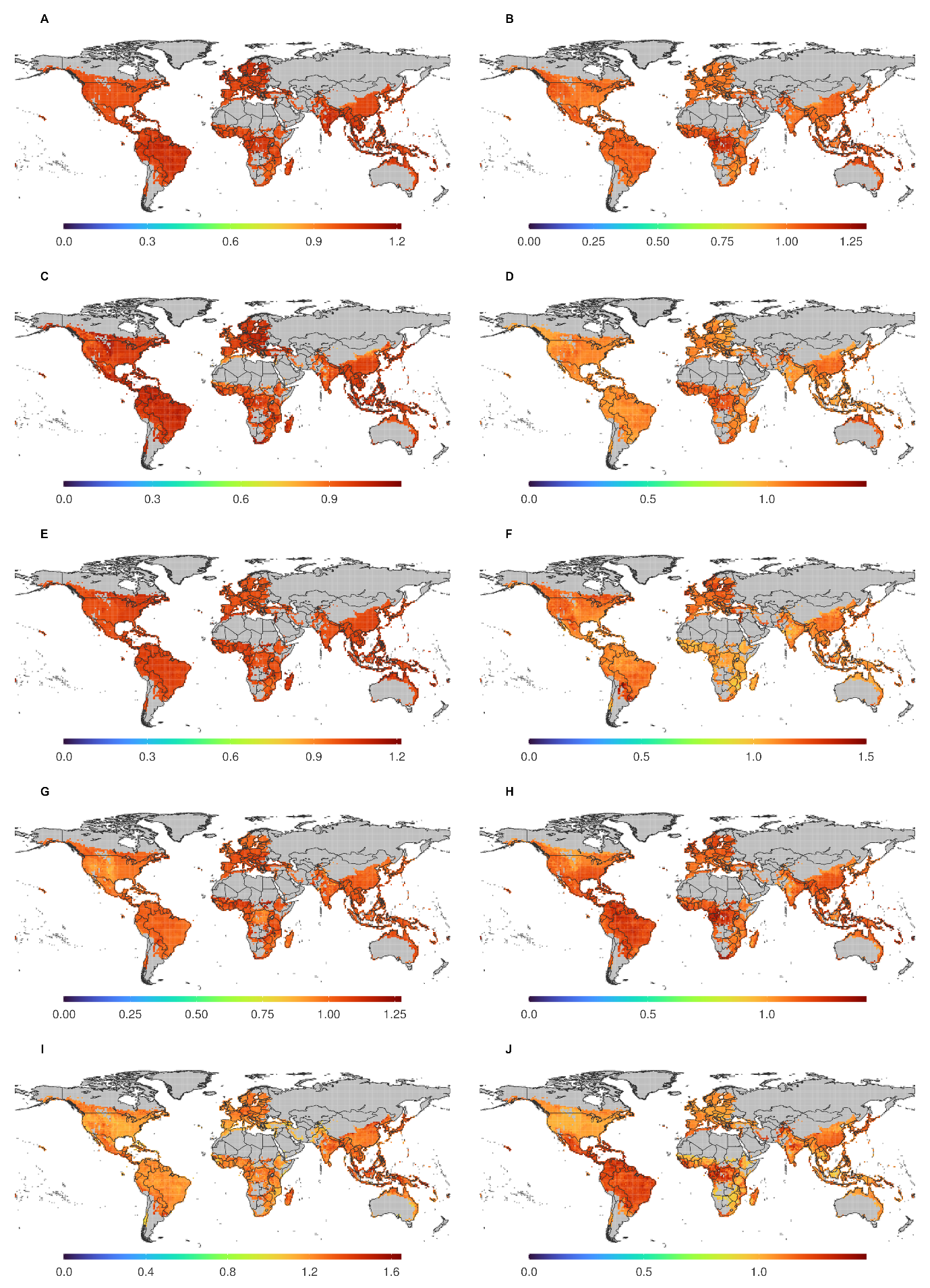

**Figure S8.** Global pattern of (A) CYP78A5, (B) CYP79A1, (C) CYP80G2, (D) CYP81E1, (E) CYP82Y1, (F) CYP83B1, (G) CYP84A4, (H) CYP85A3, (I) CYP86A1, and (J) CYP87D18

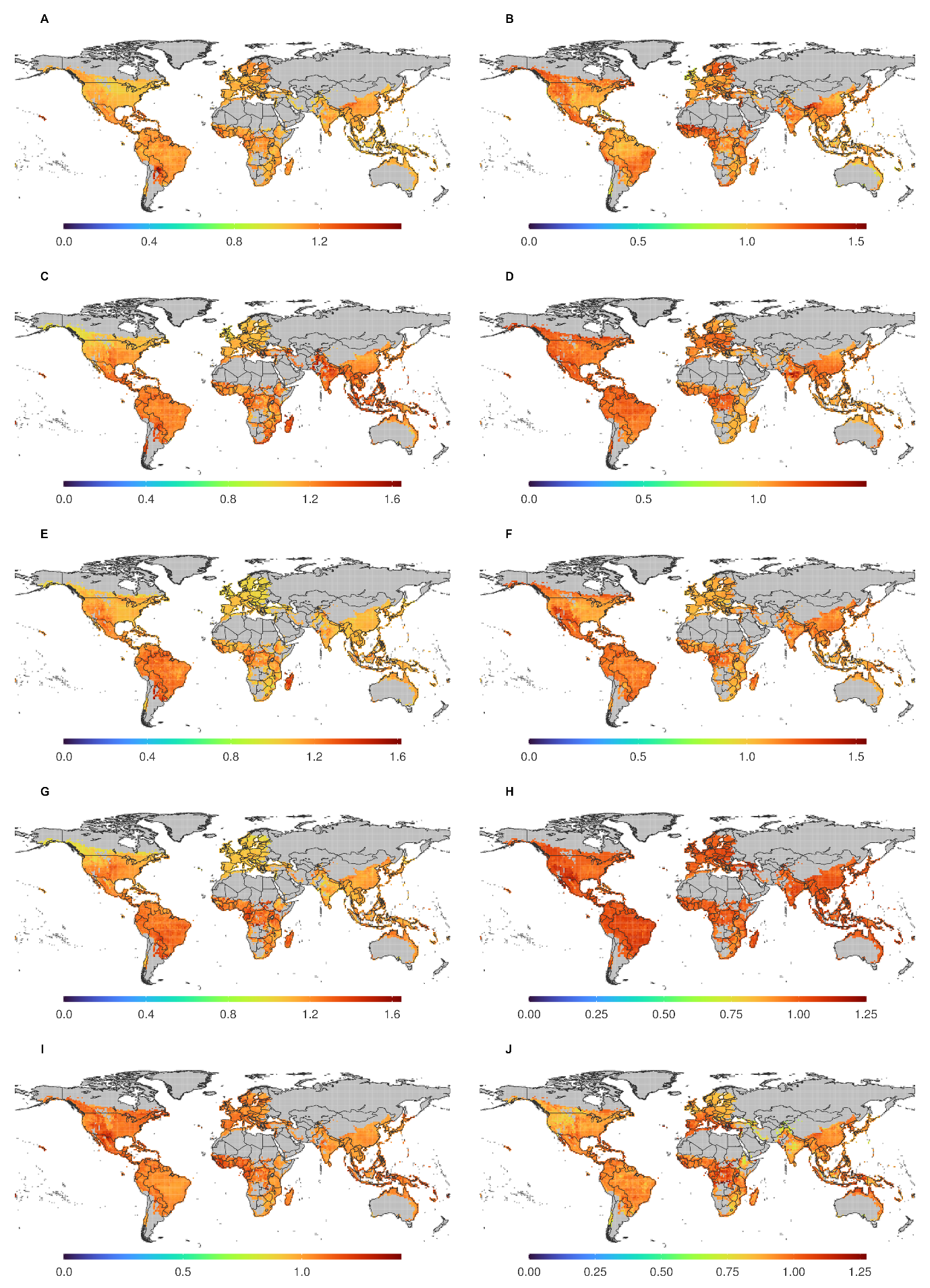

**Figure S9.** Global pattern of (A) CYP88A3, (B) CYP89A2, (C) CYP90C1, (D) CYP91A2, (E) CYP92C5, (F) CYP93C2, (G) CYP94B3, (H) CYP95, (I) CYP96A15, and (J) CYP97C1

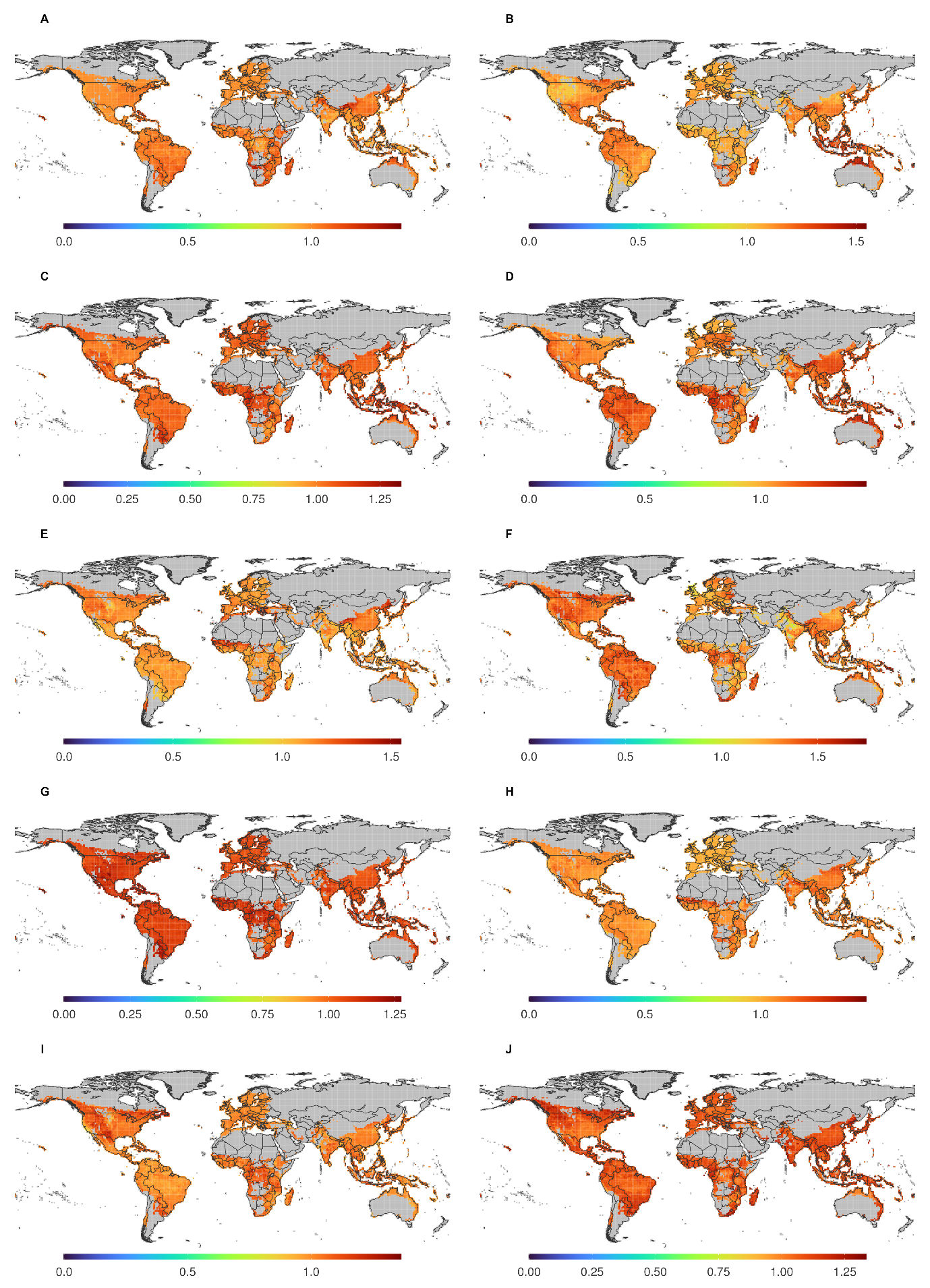

**Figure S10.** Global pattern of (A) CYP98A8, (B) CYP99A3, (C) CYP701A8, (D) CYP702A5, (E) CYP703A2, (F) CYP704B1, (G) CYP705A5, (H) CYP706A3, (I) CYP707A2, and (J) CYP708A2

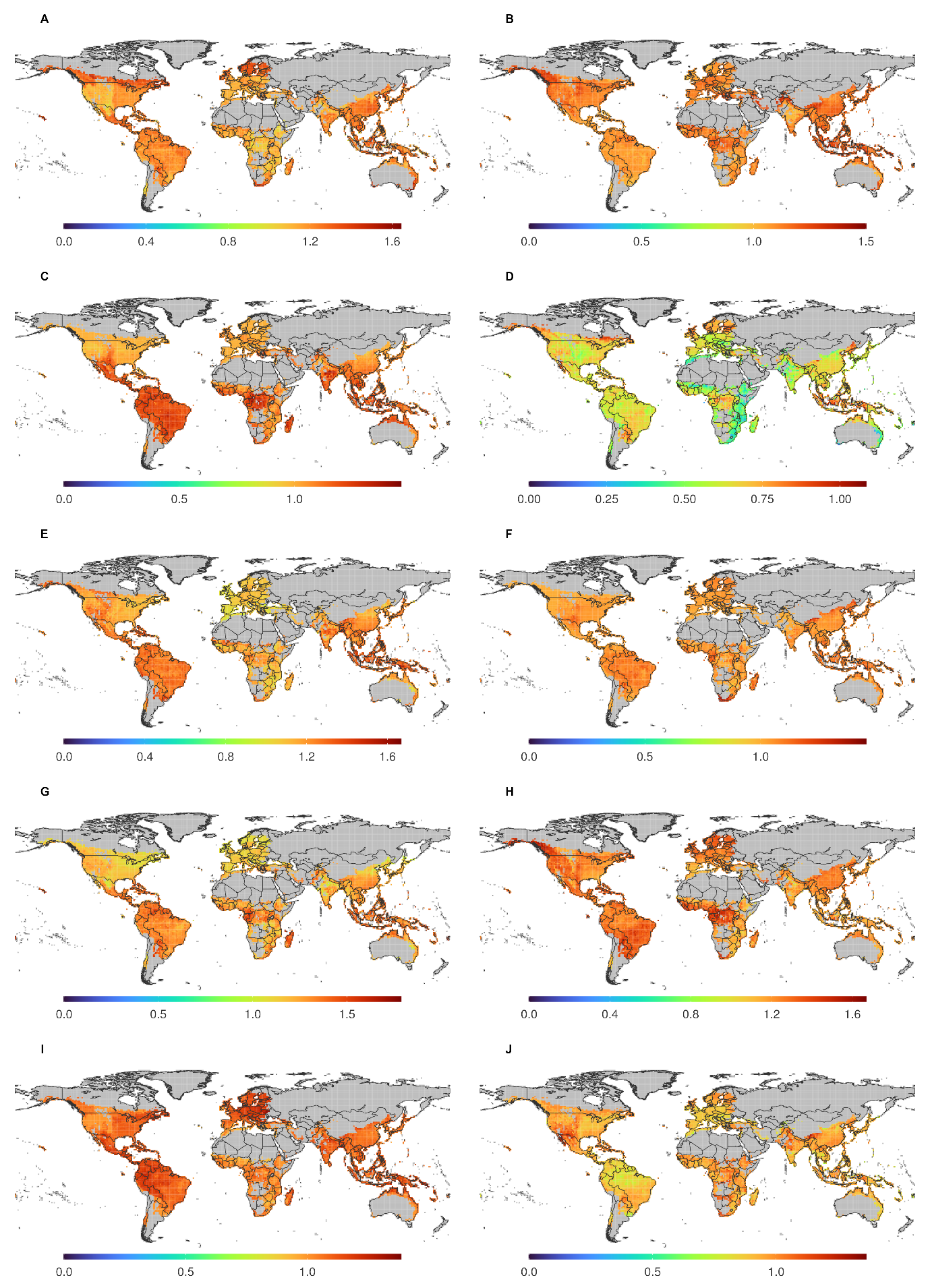

**Figure S11.** Global pattern of (A) CYP709B3, (B) CYP710A1, (C) CYP711A1, (D) CYP712A1, (E) CYP714B1, (F) CYP715A1, (G) CYP716A47, (H) CYP718B1, (I) CYP719A23, and (J) CYP720B1

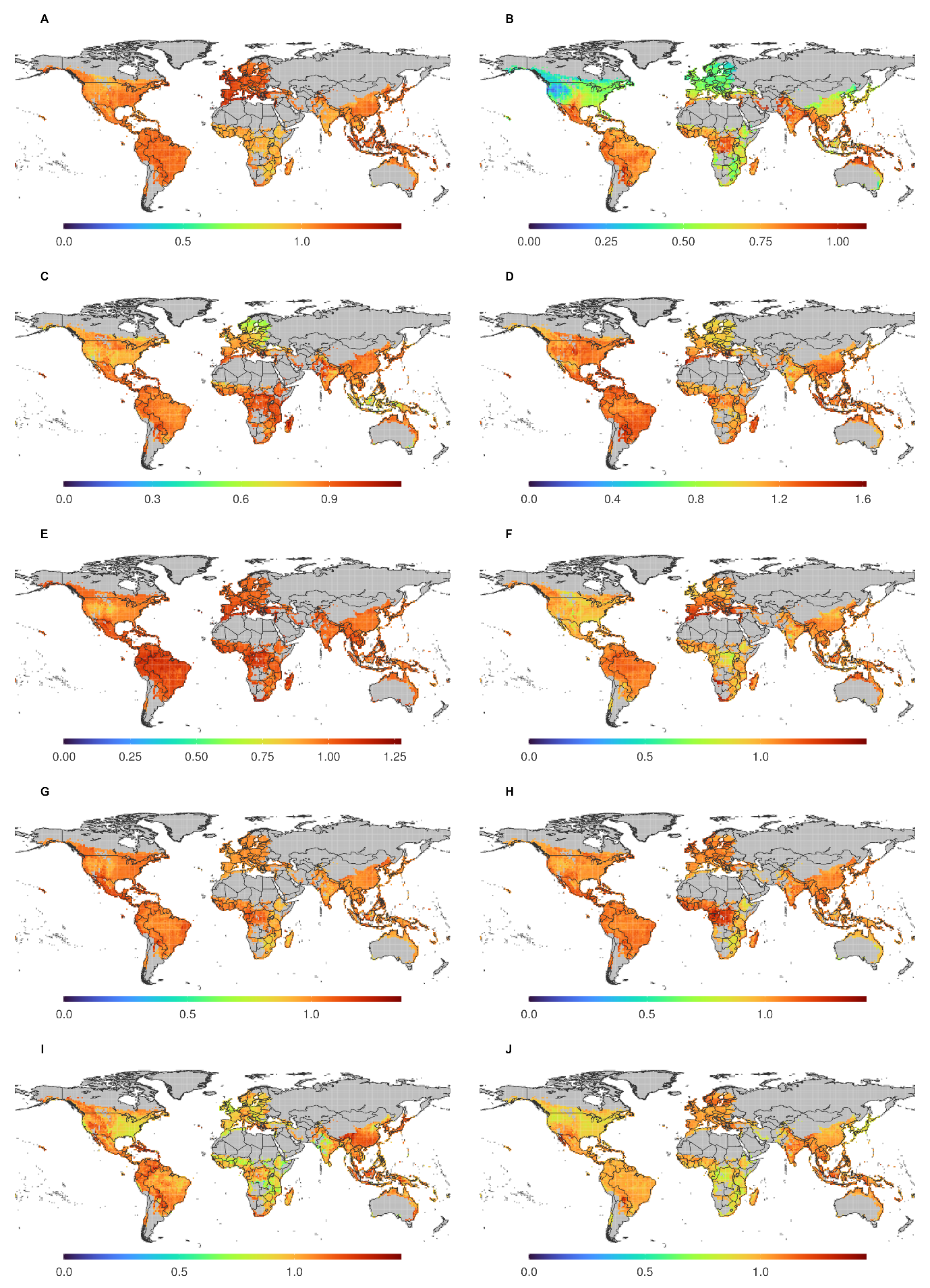

**Figure S12.** Global pattern of (A) CYP721A1, (B) CYP722A1, (C) CYP724B1, (D) CYP725A1, (E) CYP726A13, (F) CYP727C1, (G) CYP728S3, (H) CYP729B25, (I) CYP733A1, and (J) CYP734A1

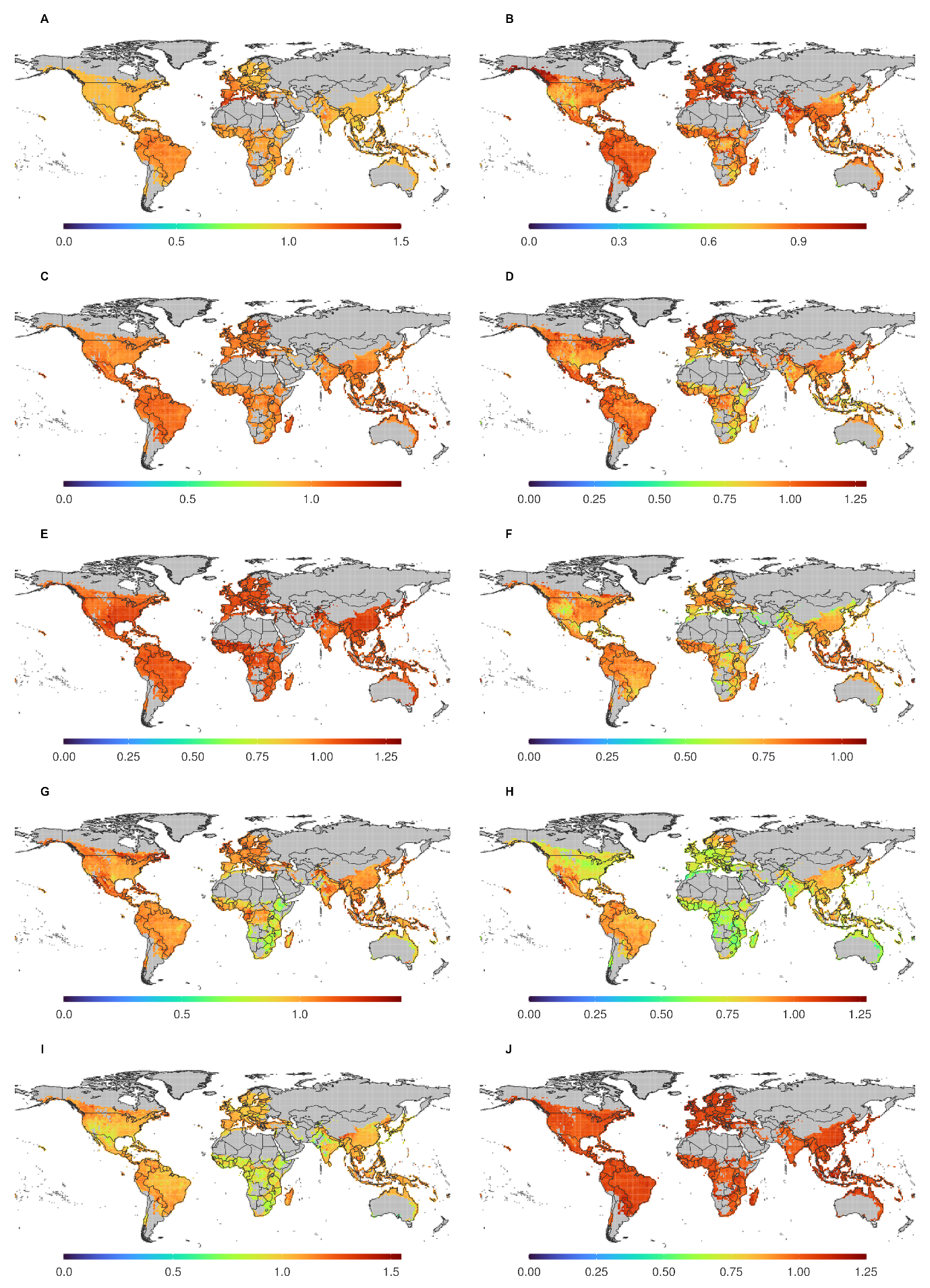

**Figure S13.** Global pattern of (A) CYP735A2, (B) CYP736A117, (C) MdPPO, (D) COX10, (E) HPT_VTE2-1, (F) ClPT1, (G) N8DT-1, (H) ABC4, (I) CHLG, and (J) PPT1

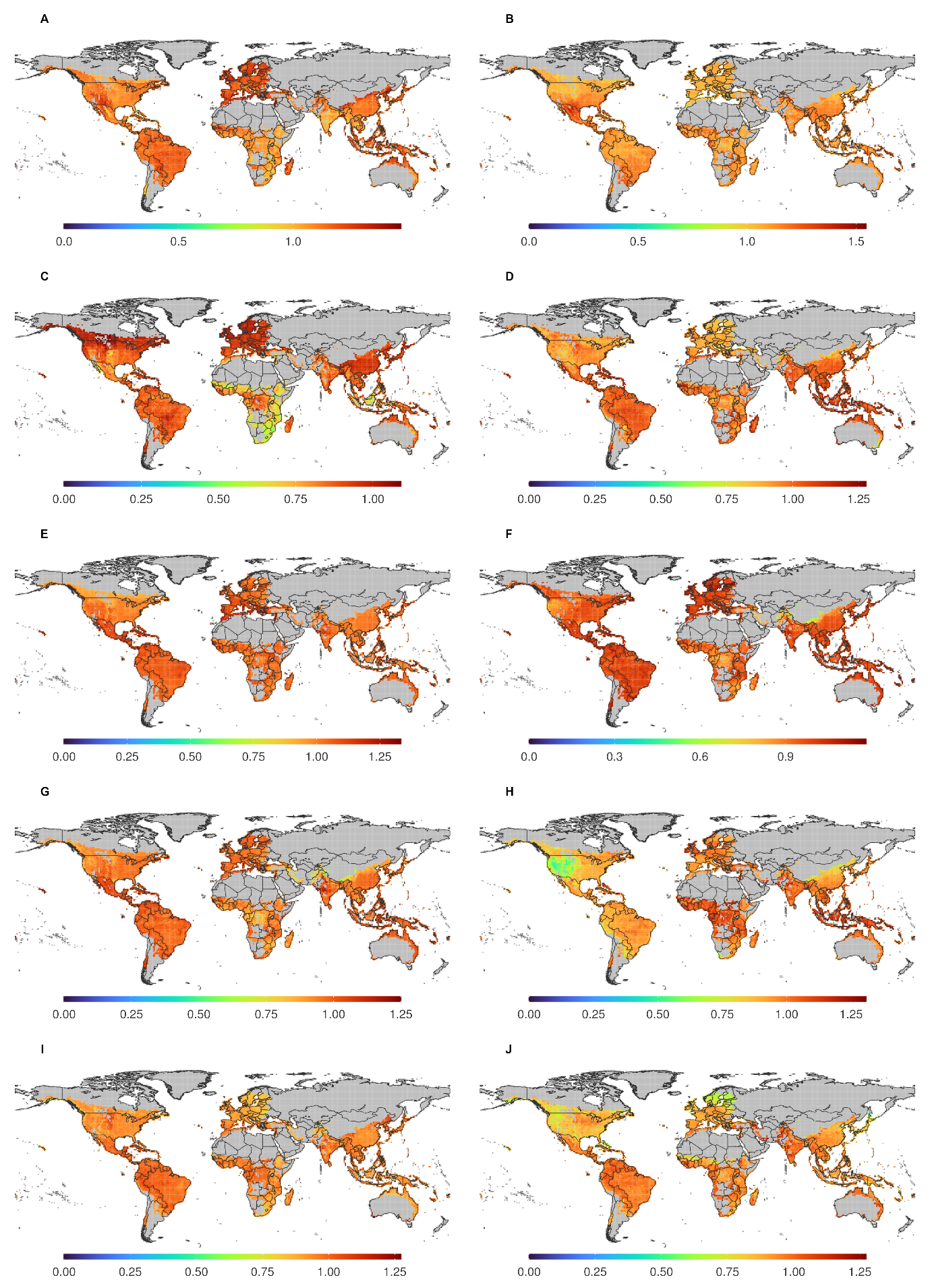

**Figure S14.** Global pattern of (A) HST, (B) PcPT, (C) AcPT1, (D) RdPT1, (E) HGGT, (F) PT1, (G) Ptpat, (H) FPT, (I) PGT-1, and (J) TPS26

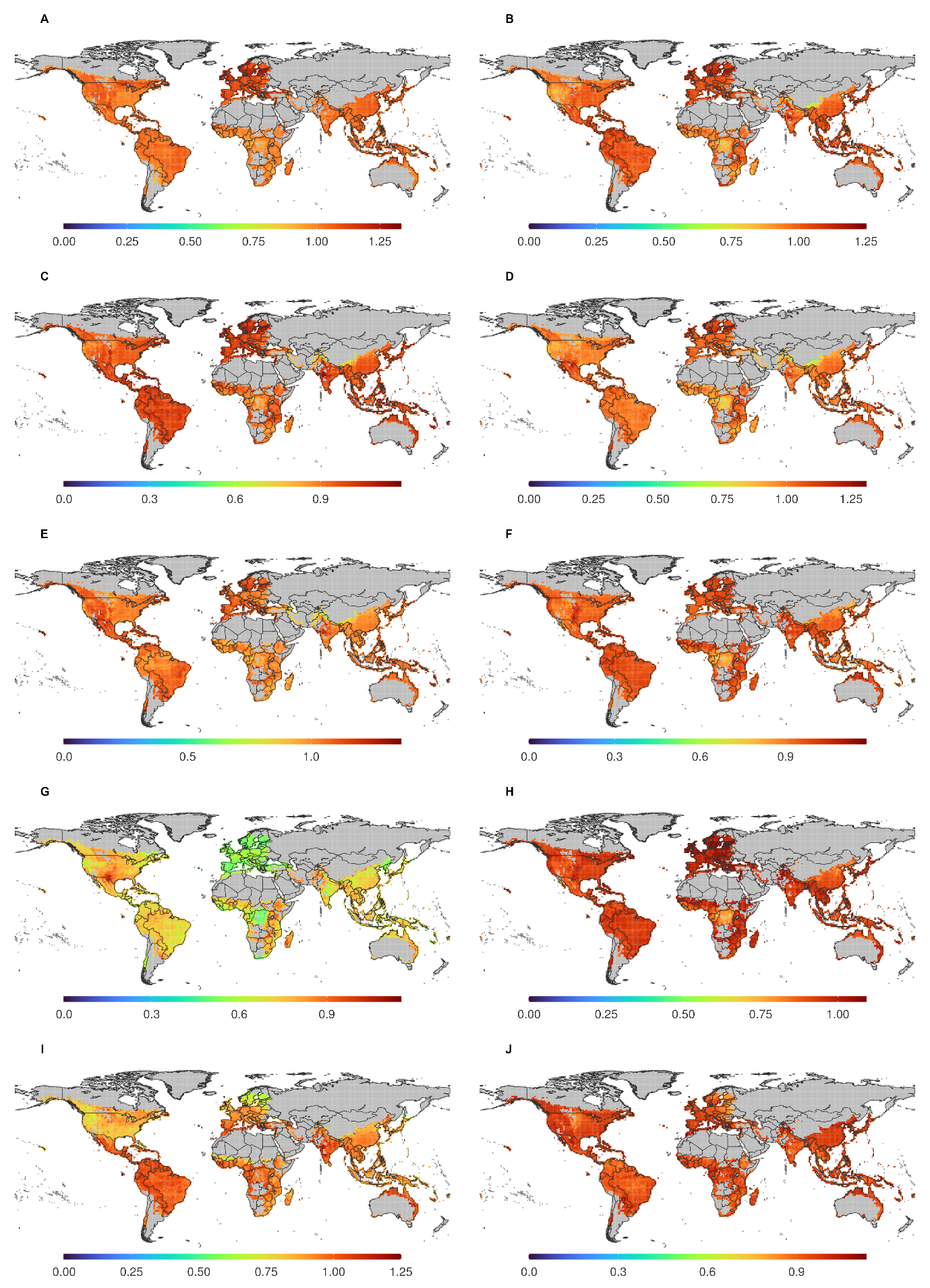

**Figure S15.** Global pattern of (A) TP20L, (B) AtTPS14, (C) TPS3, (D) TPS4, (E) TPS31, (F) TPS41, (G) smTPS4, (H) UGT71C1, (I) UGT72E2, and (J) UGT73B4

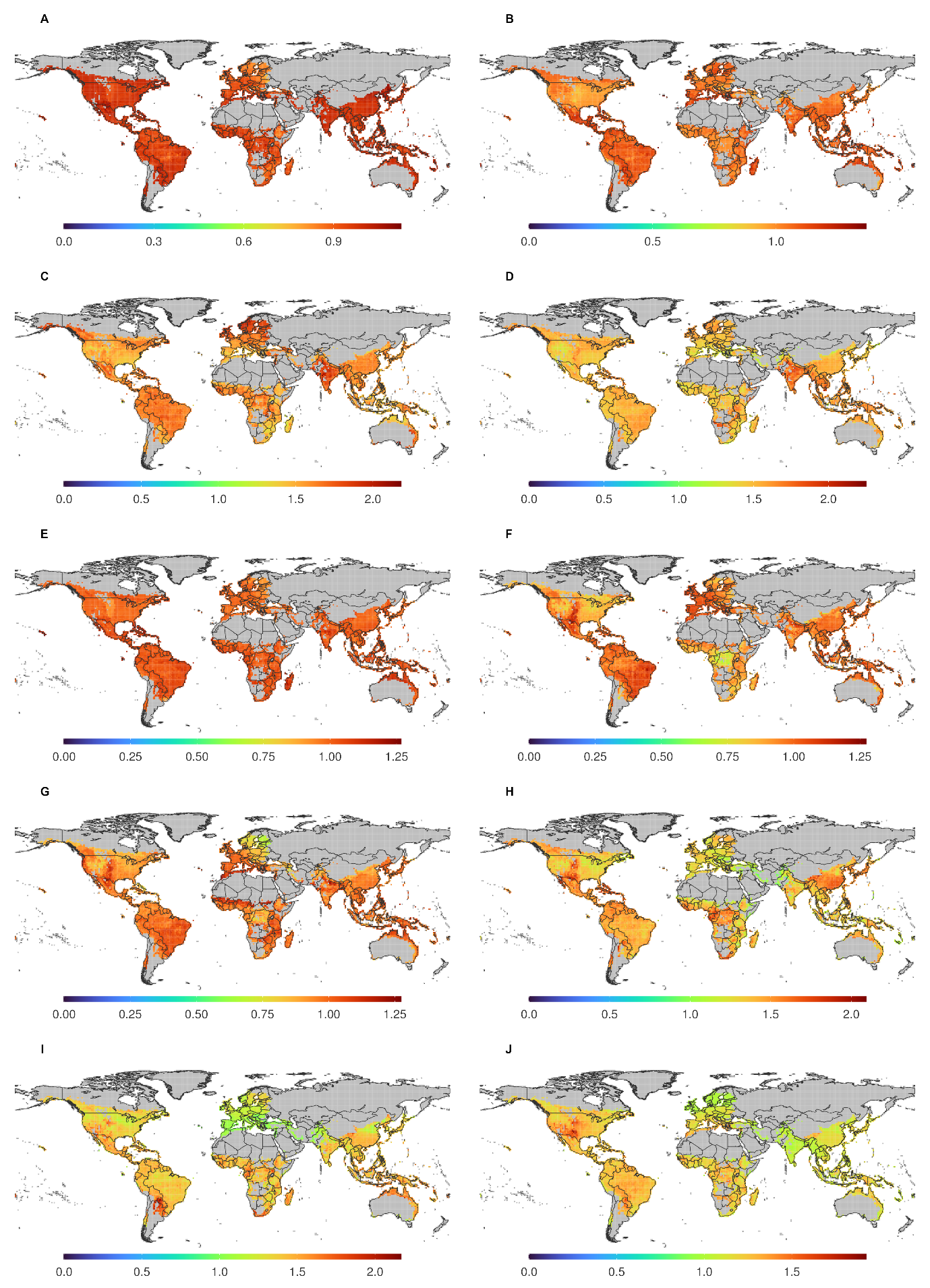

**Figure S16.** Global pattern of (A) UGT74F2, (B) UGT75B1, (C) UGT76G1, (D) UGT77B2, (E) UGT78G1, (F) UGT79B1, (G) UGT80A2, (H) UGT81A1, (I) UGT82A1, and (J) UGT83A1

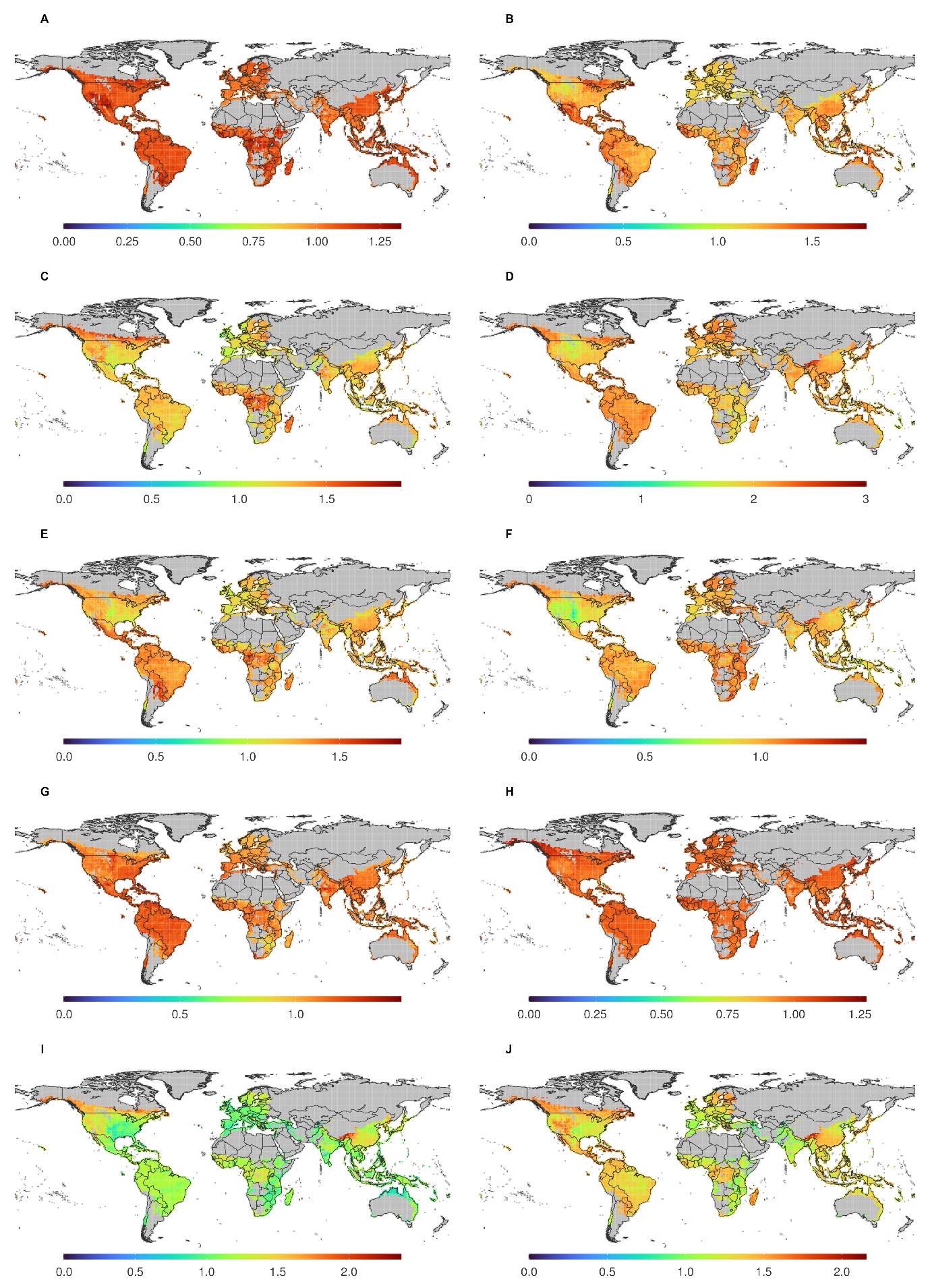

**Figure S17.** Global pattern of (A) UGT84A1, (B) UGT85B1, (C) UGT86A2, (D) UGT87A2, (E) UGT88E3, (F) UGT89C1, (G) UGT90A1, (H) UGT91D2, (I) UGT92A1, and (J) UGT93B8

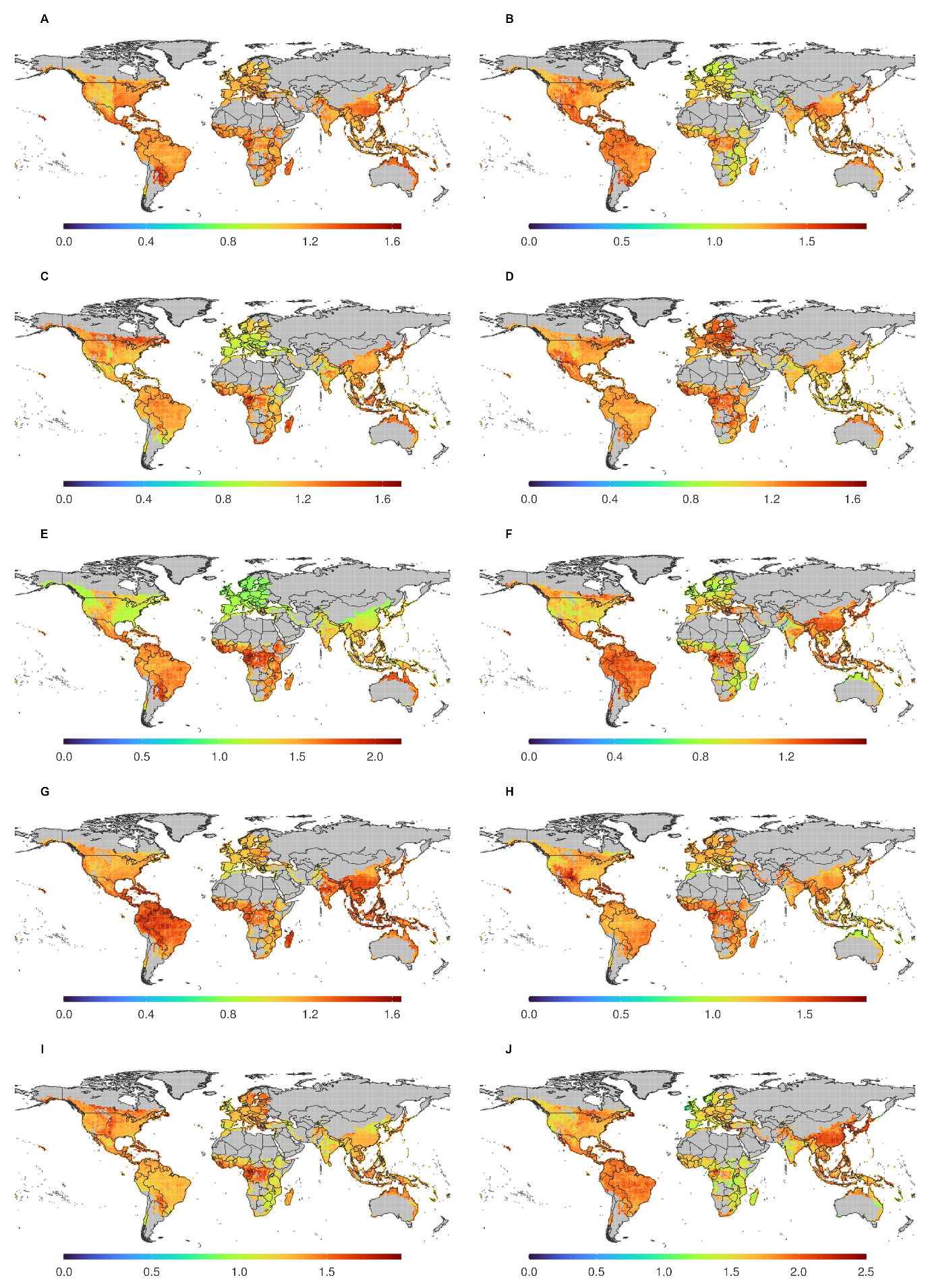

**Figure S18.** Global pattern of (A) UGT94B1, (B) UGT95B1, (C) UGT97B3, (D) UGT98B4, (E) UGT99A6, and (F) UGT100

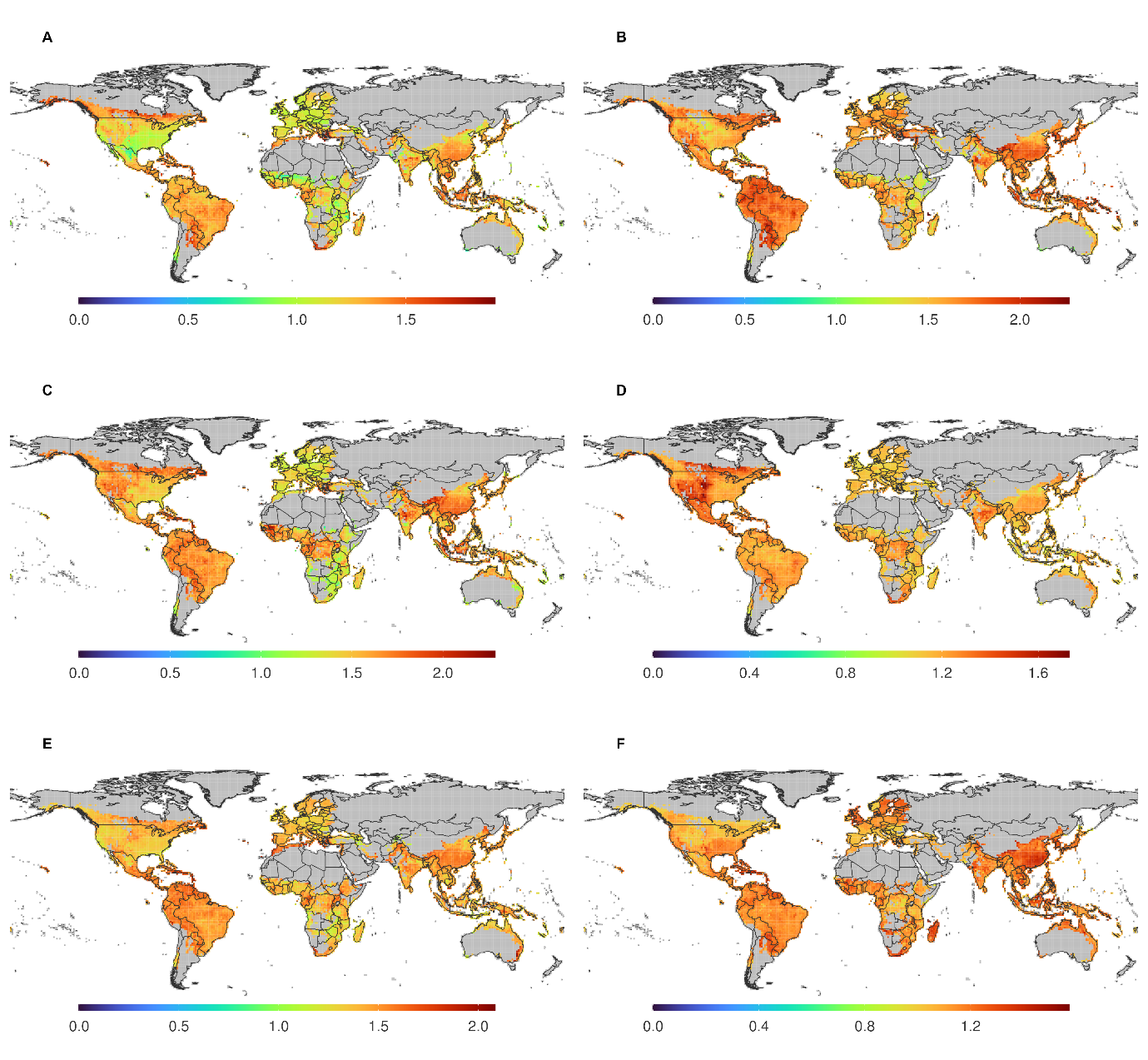

**Section 1:** De novo assembly simulation to elucidate optimal k-mer

To elucidate the optimal k-mer choice for our study, we initiated a de novo assembly simulation. We downloaded the coding sequence (CDS) and protein sequence of selected plants (Supplementary Data 2) from EnsemblePlants (release 52) as a dataset. The CDSs were later simulated to produce RNA-seq reads using Polyester v.1.36.0 with 100 reads per transcript. We performed de novo assembly following the procedure in de novo assembly for raw transcriptome data section. The performance of each assembly was assessed with simulated benchmarks and common assembly statistics, which include mean, median, longest, and N50 sizes of scaffolds or contigs, and mean coverage, which was calculated from the mapping of the assembly with its reads using BWA-MEM v.0.7.17. The simulated benchmark scores included the correct-to-incorrect ratio, accuracy, and ratio of the detected reference (appendix 3). These metrics were retrieved based on the Blastx v.2.12.0 search of each scaffold or contig in an assembly to its original protein transcripts as a reference. From the Blastx alignment, we assumed a scaffold or contig to be misassembled if its alignment result had no match to the reference transcript with an identity and sequence length threshold of 95% and 50%, respectively. Otherwise, we assumed correct assembly; the total amount of correctly assembled and misassembled was considered a true positive (TP) and a true negative (TN), respectively. Based on the alignment, we also calculated the amount of protein reference that was found and missing; the amount of missing reference was considered a false negative (FN). The ratio of correct to incorrect predictions was calculated by dividing the number of TP by the number of TN. Accuracy, on the other hand, was calculated by dividing the number of TP by the sum of TP, TN, and FN.
